# Genomic separation of Salish Sea and Pacific outer coast populations of the keystone sea star *Pisaster ochraceus*

**DOI:** 10.1101/2025.07.24.666598

**Authors:** Paige J. Duffin, Lauren M. Schiebelhut, Michael N Dawson, John P. Wares

## Abstract

Understanding dispersal and population structure in marine species is critical to forecasting ecological responses to climate change. The ochre sea star (*Pisaster ochraceus*), an intertidal keystone predator along the North American Pacific coast, has long been considered to have little spatial genetic structure due to its extended larval dispersal period and thus high potential for gene flow. Here we characterize the spatial genomic architecture of *P. ochraceus* using whole-genome sequencing (WGS) data from individuals spanning over 3,000km of coastline from Alaska to southern California. Our analyses of putatively neutral SNPs evince considerable mixing throughout the latitudinal range yet reveal substantial population structure between outer Pacific coast populations and those residing within the semi-enclosed estuary of the Salish Sea, suggesting restricted gene flow and demographic divergence between these regions. Genomic divergence is further supported by evidence of selection, with outlier loci highlighting extended regions of low diversity in the Salish Sea, consistent with recent selective sweeps and potential local adaptation to the distinct estuarine conditions. These findings are consistent with the role of oceanographic barriers or environmental heterogeneity in shaping population structure in *P. ochraceus*, challenging an earlier notion of range-wide homogeneity. By exploring neutral and adaptive genetic diversity in this keystone marine species, we provide valuable insights for its capacity to persist in a rapidly changing world.

## Introduction

The varied responses of interconnected populations, species, and communities to a rapidly changing environment is a key focus of evolutionary ecology. In marine ecosystems, there are numerous reports of widespread declines, shifting geographic distributions, and evolving interactions (Barry et al., 1995; Sunday et al., 2012; Duffin et al., 2021; Petraitis and Dudgeon, 2020; Kardos et al., 2023); the most comprehensive data typically come from economically important species (e.g. Clausen & York, 2008; Christensen et al., 2014; Lafferty et al., 2015; Benestan et al., 2016; Stewart-Sinclair et al., 2020), leaving significant gaps in our understanding of broader biodiversity (Dietrich et al., 2020). These knowledge gaps hinder our ability to predict how species and ecosystems will respond to ongoing climate change through movement and adaptation. In many cases, accurate forecasting hinges on understanding how offspring or propagules disperse and identifying critical intraspecific variation that ensures species’ persistence (Kelly et al., 2012; Bay et al., 2019; Lotterhos et al., 2019; Toczydlowski and Waller, 2019).

In marine ecosystems, a key factor shaping forecasts will be how offspring disperse (Bates et al., 2014; Wilson et al., 2016; Bashevkin et al., 2020; Pinsky et al., 2020). Larval dispersal in marine species is shaped by a complex interplay of factors, including those intrinsic to a species such as phenology and behavior of developing larvae as well as extrinsic forcing from ocean currents, upwelling, salinity and other abiotic drivers (<u>Pringle</u> et al., 2014; Bashevkin et al., 2020; Wares et al., 2021; Martins et al., 2022). Despite the challenges, identifying factors that promote or limit connectivity is fundamental to marine ecology and evolution as it provides context for the mechanisms driving local adaptation (e.g., Bohonak, 1999; Miller & Ayre, 2008; Sanford & Kelly 2011; Riginos et al., 2013; Dayan, 2020; Nielsen et al., 2020). Genetic analysis provides essential insights into dispersal processes, demographic history, and adaptive divergence amid a spatially and temporally heterogeneous seascape (e.g., Benestan et al., 2016; Liggins et al., 2020; Papa et al., 2022; Wadgymar et al., 2022). While initially, genetic patterns clarified the ways in which intrinsic and extrinsic factors may affect patterns of gene flow within species (Palumbi 1994, Wares et al 2001, Dawson et al 2014), ever-advancing molecular tools have revealed a vastly increased resolution on the spatial genetic structure of marine species, including those with high dispersal capacity and few clear barriers to movement (e.g., Hess et al., 2013; Berg et al., 2015; Jeffrey et al., 2017; Hoey et al., 2018).

Through this increased resolution, genomic analyses now allow us to more clearly isolate the effects of dispersal from demographic and adaptive patterns. This is of key importance when studying ecologically significant organisms that may vary in response to a changing environment, and understanding the trajectory of these species guides our understanding of whole communities (Power et al., 1996). We are particularly interested in broadly-distributed species that, in studying patterns of genomic diversity, guide us on the mechanisms leading to biogeographic patterns as well as evolutionary response to key environmental gradients (Dawson, 2001; Wares et al., 2001; Ewers-Saucedo et al., 2015). Adding population genomic data to such systems enables transformative capacity to resolve spatial and evolutionary dynamics, in particular through recognizing genomic “footprints of local adaptation” (Wilder et al 2020).

The ochre sea star (*Pisaster ochraceus*) is a conspicuous and ecologically significant shallow-water sea star of the North American Pacific Coast. As a keystone predator, *P. ochraceus* plays a vital role in maintaining biodiversity within rocky intertidal ecosystems (Paine, 1966, 1969, 1974; Harley, 2003; Blanchette et al., 2005; Robles et al., 2009; Pearse et al., 2010; Lawrence, 2013). These broadcast-spawning asteroids have a lengthy dispersal stage facilitated by feeding larvae that can survive in a pelagic state for months (Strathmann, 1978, Sanford & Menge, 2007) but have a typical metamorphosis to juvenile recruitment around 6-8 weeks (Pia et al., 2012).

Along with these dispersal traits, prior genetic analyses have suggested very little spatial pattern of diversity (Stickle et al., 1992; Harley et al., 2006; Frontana-Uribe et al., 2008; Marko et al., 2010; Schiebelhut et al., 2023), yet indications of metapopulation dynamics leaving transient signatures of ‘chaotic genetic patchiness’ (Johnson & Black, 1982; see Schiebelhut et al., 2022a). With a distribution spanning over 4800 km, ochre star populations experience a vast array of environmental conditions; for example, monthly sea surface temperatures near the southern distributional limit in Baja California, Mexico, are nearly 10°C higher than at the northern limit in Prince William Sound, Alaska, USA (see Supplement S1 and Dawson et al., 2024).

In addition to a suite of factors that vary latitudinally, ochre stars are found within multiple distinct coastal habitats, from the exposed open seaboard to sheltered estuaries and fjords, such as the Salish Sea (Figure 1). This glacially-formed estuarine basin is expansive (17,803 km^2^; Flower, 2020) and contains a unique ecosystem, circulation, and stratification (e.g., Ebbesmeyer et al., 1984; Sutherland et al., 2011; Khangaonkar et al., 2019; Barth et al., 2019; MacCready et al., 2020; Sobocinski, 2021). The ecosystem of the Salish Sea has historically supported an abundance of *P. ochraceus*, making it a key locale for ochre star research. The Salish Sea receives seasonal freshwater input primarily from the Fraser River, with smaller contributions from surrounding watersheds (Sobocinski, 2021), yet maintains salinity similar to the outer coast due to continuous marine inflow through several large straits. Though technically an estuary, it remains more saline than comparable systems like San Francisco Bay, where *P. ochraceus* is absent. Despite apparent connectivity with the Pacific, the Salish Sea hosts striking faunal diversity and endemicity. Even highly mobile species, such as killer whales (*Orcinus orca*), maintain genetically and behaviorally distinct resident populations (Barrett-Lennard & Ellis, 2001; Morin et al., 2024). This unique configuration sets up key contrasts between the biodiversity of the Salish Sea and the outer coast.

**Figure 1.**
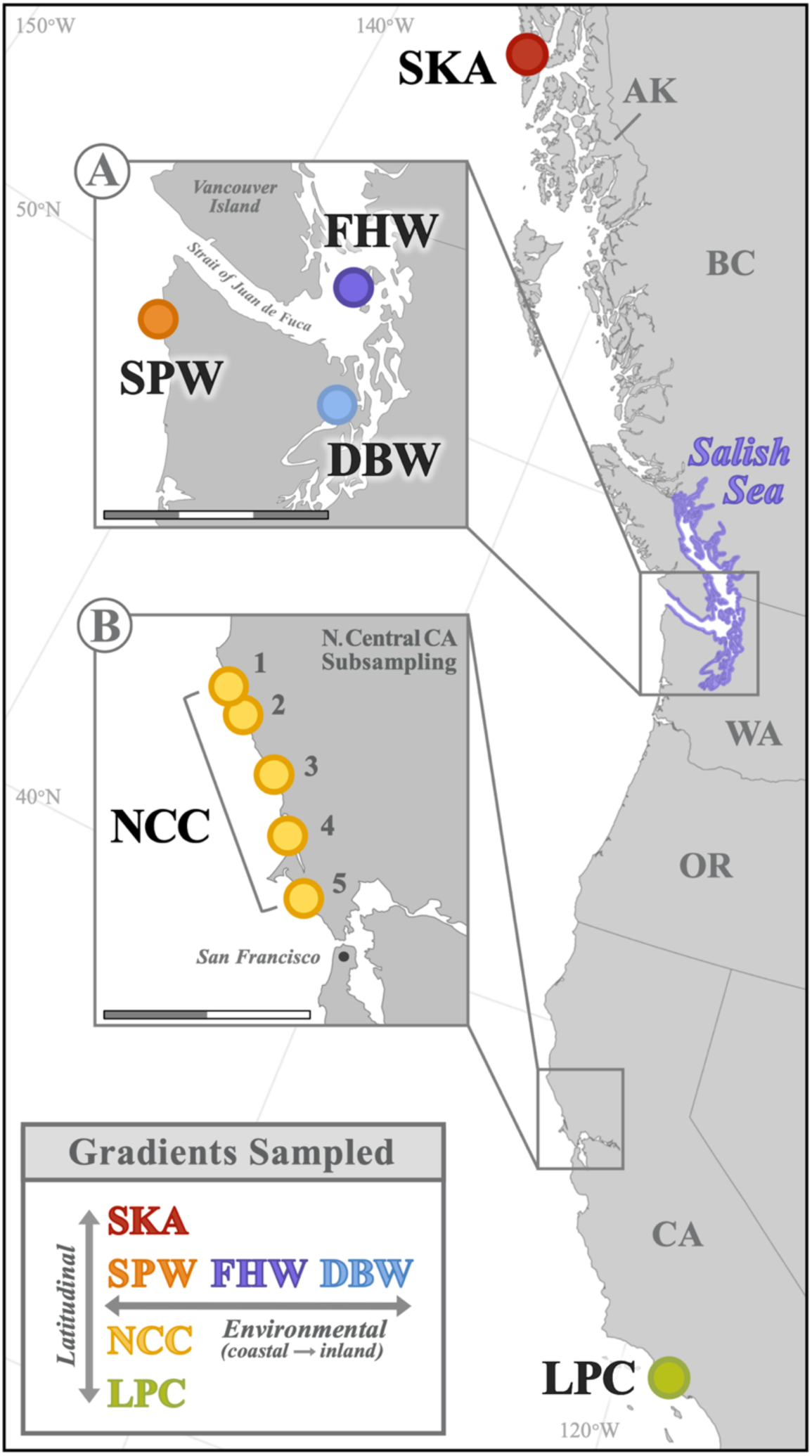
**Sampling locations included in this study**. The six major sampling regions were: <u>S</u>it<u>k</u>a, <u>A</u>laska (SKA, n=7); <u>S</u>okol <u>P</u>oint, <u>W</u>ashington (SPW, n=8); <u>F</u>riday <u>H</u>arbor, <u>W</u>ashington (FHW, n=10); <u>D</u>abob <u>B</u>ay, <u>W</u>ashington (DBW, n=2); five subsampled regions in <u>N</u>orth <u>C</u>entral <u>C</u>alifornia (NCC, n=60) and <u>L</u>echuza <u>P</u>oint, <u>C</u>alifornia (LPC, n=8). One scale bar unit (alternating black and white) is equal to 50 km. Regional and inset maps created using SimpleMappr (Shorthouse, 2010) using the North American Lambert projection and default longitude of natural origin −96. See Table 1 for additional sampling location details.

**Table 1.** Detailed information for the sampling locations included in this study.

| Sampling Location |  | n | Lat. | Long. | Date Sampled |
| --- | --- | --- | --- | --- | --- |
| **SKA** | Sitka, AK | 7 | 57.075 | -135.372 | 02/2018 |
| **SPW** | Sokol Point, WA | 8 | 47.947 | -124.662 | 05/2018 |
| **FHW** | Friday Harbor, WA | 10 | 48.551 | -123.006 | 06/2016 |
| **DBW** | Dabob Bay, WA | 2 | 47.77 | -122.8 | 07/2004 |
| **NCC** | North Central CA | 60 | ---- | ---- | 12/2014<br>through<br>06/2015 |
| **NCC<sub>1</sub>** | Serenisea | 12 | 38.798 | -123.573 |  |
| **NCC<sub>2</sub>** | Sculpture Point | 12 | 38.7 | -123.443 |  |
| **NCC<sub>3</sub>** | Twin Coves | 12 | 38.459 | -123.146 |  |
| **NCC<sub>4</sub>** | McClures Beach | 12 | 38.182 | -122.966 |  |
| **NCC<sub>5</sub>** | Palomarin | 12 | 37.931 | -122.75 |  |
| **LPC** | Lechuza Point, CA | 8 | 34.034 | -118.86 | 01/2018 |

Our focus on the biology of *P. ochraceus* across these distinct environmental transitions follows a series of long-standing questions about striking color polymorphism patterns and gradients (Harley et al., 2006; Raimondi et al., 2007), and about how *P. ochraceus* maintains physiological and phenotypic plasticity across diverse habitats. Here we evaluate whether subtle genetic structuring exists below previous detection thresholds using single marker, reduced representation, or more geographically restricted approaches. Whole-genome sequencing provides the opportunity to assess these patterns with orders of magnitude greater resolution.

Leveraging the recently updated *P. ochraceus* genome (Ruiz-Ramos et al., 2020; Hu et al., *in prep*), we conducted whole-genome resequencing of 65 individuals spanning more than 3000 km of coastline and a major environmental transition, from the open Pacific to the brackish Salish Sea (Figure 1). Given growing concern for the persistence of this iconic species—largely due to a range-wide mass mortality event that affected many species of sea stars (Eisenlord et al., 2016; Kohl et al., 2016; Menge et al., 2016; Aquino et al., 2021; Oulhen et al., 2022; Dawson et al., 2023), our aim is to resolve fine-scale genetic structure and assess the role of both broad-scale geographic variation and localized environmental factors in shaping patterns of genetic diversity that may guide our understanding of the persistence of *P. ochraceus*.

## Materials and Methods

### Sample Collection

A total of 65 adult *Pisaster ochraceus* individuals were sampled intertidally from 10 sites spanning nearly 3000 km of coastline and representing most of the species’ range (Table1, Figure 1). The distribution of these sites is partly associated with other studies on genomic diversity of *P. ochraceus*, which thus fall into four latitudinal groups: [1] our northernmost site at Sandy Beach, in Sitka, AK (SKA); [2] the outer coast of Washington state (Sokol Point, SPW); Friday Harbor, WA (FHW), located in the San Juan Islands of the Salish Sea; Dabob Bay, WA (DBW), a sheltered saltmarsh bay branching off Hood Canal in Puget Sound (and the greater Salish Sea); [3] five subsampled locations along an ∼120 km transect in north central California (NCC_1_ through NCC_5_, see Table 1 for site names); [4] the southernmost site, Lechuza Point, CA (LPC), just south of Point Conception, in a widely-recognized biogeographic transition zone (e.g., Wares et al., 2001; Blanchette et al., 2007; Briggs & Bowen, 2012, 2013; reviewed in Dawson, 2001) (Figure 1). In addition to the latitudinal sampling, our study design captures points on both the outer coast (OC) and sheltered intracoastal waterways in the Salish Sea (SS) which includes Puget Sound.

Tissue collection consisted of nonlethal amputation of 10-30 tube feet, which were immediately placed in 95% ethanol (9/10 sites) or DMSO buffer (DBW only). DNA isolation protocols varied across samples, with those from SKA, SPW, and LPC following Schiebelhut et al. (2022); those from FHW and DBW were isolated as described in Chandler & Wares (2017).

DNA sequencing was carried out in two sets. Purified genomic DNA from NCC samples were submitted to the DNA Technologies & Expression Analysis Core at the University of California, Davis, USA for RNase treatment, library preparation, quality control, and sequencing. Library traces were assessed using a LabChip GX prior to pooling. Libraries were PE150 sequenced on an Illumina NovaSeq S4 – S4 flowcell lane.

Genomic DNA from samples collected at the remaining five sites (SKA, SPW, FHW, DBW, and LPC) were prepped for blunt-end ligation using End-It^TM^ DNA End-Repair Kit (Epicentre Biotechnologies) according to the manufacturer’s instructions. DNA was cleaned using AMPure beads (Beckman Coulter) to select fragments of 100 bp or higher, and prepped for Illumina sequencing as in Ji et al. (2018).

### Data Processing

#### Sequence Processing / SNP Calling

Raw paired-end reads were trimmed for quality using Trim Galore! (Krueger, 2015), and sequence quality was assessed with FastQC (Andrews, 2010). Reads were mapped to the *P. ochraceus* reference genome (version 2, Hu et al., *in prep*) using BWA MEM (Li & Durbin, 2009). Standard post-processing steps followed Genome Analysis Toolkit (GATK) best practices (Broad Institute, DePristo et al., 2011), including duplicate removal, sorting, and mate-pair correction using Picard and SAMtools (Li et al., 2009).

Variants were called using BCFtools (Li et al., 2009), and the resulting variant call format (VCF) file was used to generate a consensus sequence. VCFtools (Danecek et al., 2011) was used to calculate summary statistics before quality filtering. Full computational details and commands are provided in Supplement S2.

#### Quality Filtering

Variants were first filtered using VCFtools by removing the following: (**1**) indels (--remove-indels); (**2**) SNPs missing data in > 20% of individuals (--max-missing 0.8); (**3**) SNPs with QUAL base quality scores below 200 (--minQ 200). Filtering continued in R v4.02 (R Core Team, 2020, version v4.02 for all R-based analyses) using RStudio (RStudio Team, 2020), where additional SNPs were removed as follows: (**4**) SNPs with depth (DP) below 200 or above 2500; (**5**) SNPs with MQ (mapping quality) values below 40. Finally, VCFtools was used again to remove the following SNPs: (**6**) SNPs missing data in > 10% of individuals (--max-missing 0.9); (**7**) SNPs with a minor allele frequency (MAF) below 5% (--maf 0.05). Supplement S2 provides the number of SNPs removed at each quality-filtering step to generate a final SNP panel of 11,701,389 high quality SNPs along with additional details on bioinformatic processing.

#### Linkage Disequilibrium

We calculated linkage disequilibrium separately for each of the main 22 chromosomes using VCFtools (v 0.1.16) (Danecek et al., 2011) and the following generalized code: vcftools --gzvcf quality.filt_SNPs.vcf.gz --geno-r2 --ld-window-bp 10 --out LD_calc_10.bp.windows. See Supplement S3 for additional details.

#### Data Partitioning

We performed principal component analysis (PCA) using the R package PCAdapt (v4.3.3) to investigate population structure across the full SNP dataset. SNPs were parsed by chromosome and converted from vcf to bed file format to conduct exploratory PCAs by setting K = 20 to examine the major axes of genetic variation. Based on patterns revealed in PCAdapt and prior findings of chromosomal inversions in related species (DeBiasse et al., 2022), we excluded chromosome 9 from downstream analyses due to its disproportionate influence on PC2 (results not shown).

Putative outliers were identified and removed to generate our neutral SNP panel based on initial PCA loadings using PCAdapt (v 4.3.5) (Luu et al., 2017) and OUTFLANK (v 0.2) (Whitlock & Lotterhos, 2017) set to identify a liberal number of outliers with alpha=0.1. Finally, we again ran OUTFLANK with populations defined according to their geographic locations (alpha=0.1).

Following outlier removal, we thinned SNPs occurring within 1,500 bps of one another. Then, guided by our visualization of LD by chromosome (Supplement S3), we identified and subsequently pruned SNPs still in LD (R^2^>0.2).

Finally, an outer coast-only dataset was generated by retaining only OC-collected individuals after step 5 of the Quality Filtering pipeline, followed by VCFtools filtering to remove SNPs with a high proportion of missing data (--max-missing 0.9), rare SNPs (--maf 0.05), and SNPs that were not biallelic. This OC-only panel, which included both putatively adaptive and neutral SNPs, was used to explore fine-scale population structure along the outer coast via a PCA in R using the PCAdapt package (Luu et al., 2017).

### Overall Genetic Variation and Differentiation

Global F_ST_, D_XY_, and π values were calculated on the quality-filtered all-sites vcf for each sampling location (for π) or pair of sampling locations (F_ST_ and D_XY_) using pixy (v 1.2.7.beta1) (Korunes & Samuk, 2021) in 15kb windows, then averaged across all windows along the genome. Specifically, estimates of π reflected the average per site nucleotide diversity, weighted by the number of genotyped samples at that site in that population for each 15kb window (Korunes & Samuk, 2021). Nucleotide divergence between populations (D_XY_) was calculated as the average nucleotide divergence at each site within each genomic window (Korunes & Samuk, 2021).

### Population Structure Analysis on Neutral SNPs

Principal component analyses (PCAs) were performed using PCAdapt (Luu et al., 2017) in R, with subsequent scree plots used to ensure the results were well-represented by the first two PCs (i.e., the eigenvalues of factors). We also assessed population structure across individuals using DAPC (Jombart et al., 2010) using the adegenet package (Jombart, 2008) in combination with package vcfR (v 1.12.0, Knaus & Grünwald, 2017).

Ancestry probabilities across individual genotypes were estimated with sNMF (sparse nonnegative matrix factorization, Frichot et al., 2014) through the R package LEA (v 2.20, Frichot & Francois, 2015). We tested three repetitions each of K=1 through K=12, calculated cross entropy, and qualitatively assessed the congruence of patterns across repetitions. While the cross-entropy value was lowest for K=1 and increased steadily through K=12, we noted the congruence of patterns across all repetitions of K=2 and chose to interpret this as the best value for reasons further detailed in the corresponding section of our results.

Individual ancestries were also estimated using ADMIXTURE 1.3.0 (Alexander et al., 2009). We tested three repetitions each of K=1 through K=12, calculated cross entropy, and qualitatively assessed the congruence of patterns across repetitions as described above, using the generalized command admixture -s [random seed value] vcf_converted_2_bed.bed [chosen K value].

Finally, we constructed a cladogram by first calculating a Nei’s distance matrix-based phylogenetic tree in dartR (Gruber et al., 2018). After determining bootstrap support for the Salish Sea clade, the information was transferred as a NEXUS file into FigTree (Rambaut, 2010), where the final structure was converted into a cladogram for visual representation.

### Assessing IBD Along the Outer Coast

IBD was assessed among outer coast individuals isolated from the neutral SNP panel using the Mantel test function (based on Pearson’s product-moment correlation) in the adegenet package (Jombart, 2008) based on 999 permutations of the data under the null hypothesis that there is no spatial structure present.

### Mitochondrial Analyses

We analyzed mitochondrial genome variation in *P. ochraceus* to compare our findings with previous studies on population structure. Harley et al. (2006) sequenced a 543-bp portion of mtDNA within the cytochrome c oxidase 1 (COI) gene to assess variation among 345 ochre stars across a region that largely overlaps with our sampling area, finding minimal population structure.

To facilitate direct comparison, we performed a subset of the same population structure analyses on quality-filtered SNPs mapped to the *P. ochraceus* mitochondrion (scaffold ID: ‘Scaffold_122 1_contigs length _16217’). Due to the overall low number of SNPs (171 bases) in the 16,609 bp mitochondrial genome, we did not discriminate between neutral and non-neutral sites.

### Outlier Detection / Analyses

#### Selective Sweep and Meadow Plots

Estimates of π and F_ST_ were calculated in 15 kb windows using pixy as previously described. The interval of 15 kb windows was selected based on the *P. ochraceus* genome size of 489.374 Mb in conjunction with a literature review of window size selection in other published studies as a function of their genome size (Duffin 2024). For each window, both F_ST_ and the π ratio between populations under comparison was calculated to identify regions likely involved in selective sweeps or other driving molecular mechanisms. Outliers were defined for each pairwise comparison independently as the 1% right tail (top 1%) of the empirical F_ST_ distribution simultaneously representing either the left or right 1% of empirical π ratio values. This threshold is conservative among other recent work utilizing F_ST_ and π ratios to define putative selective sweep windows; some use 1% (e.g., Zhang et al., 2020; Chen et al., 2022), while many use 5% (e.g., Li et al., 2020; Feng et al., 2021; Guo et al., 2021; Nannan et al., 2022; Sang et al., 2022). A custom R script (R v4.02) was written to generate meadow plots; representative code is available (see *Data Availability*). Stretches of contiguous outlier windows were quantified using another custom R script, allowing for up to three non-outlier windows amidst these high-Fst regions.

## Results

### Whole Genome Sequencing

Raw sequence data are archived at NCBI (SRA BioProject ID PRJNA1117092). After initial read quality filtering, adapter trimming, and mapping to the *P. ochraceus* genome, we generated a vcf dataset of 26,340,914 SNPs. After quality filtering (Supplement Table S2.1), our final dataset consisted of 11,701,389 high-quality bi-allelic single nucleotide polymorphisms (SNPs). Mean read depth across all individuals surpassed our target of 30× at 33.012 +/- 9.975 and ranged from 54.107 (star collected from LPC) to 14.538 (star collected from NCC).

#### Neutral population structure reveals divergence between outer coast and Salish Sea stars

The putatively neutral dataset included 141,625 SNPs, after removing 137,219 putatively adaptive loci from the quality-filtered full SNP dataset of the 22 main chromosomes (minus chromosome 9, see *PCA* section below for explanation) and further thinning based on linkage disequilibrium (Supplement S2 and S3 describe the data and linkage disequilibrium used for thinning).

#### Genetic Variation and Differentiation Among Sampling Locations

##### Descriptive Statistics

Average per site nucleotide diversity (π) ranged from 0.00850 (FHW) to 0.00894 (LPC). Generally, genome-wide π estimates were lowest in Salish Sea sampling locations (FHW and DBW) and higher in sites along the outer coast of North America (especially in SKA, SPW, and LPC). Our estimate of π within the densely-sampled outer coast site of NCC (n = 30) fell somewhere in the middle at 0.00884 (Supplement Table S4.1). These values are comparable to recent high-resolution estimates of π made for *Acanthaster solaris* (Popovic et al., 2024).

##### Principal Component Analysis (PCA)

PCA results on the full SNP panel revealed two prominent lines of divergence: the first along PC1, splitting sampling locations based on their geography, either in the Salish Sea (FHW and DBW) or the outer coast (SKA, SPW, NCC, and LPC). The second axis, PC2, separated individuals arbitrarily. The clusters along PC2 are driven entirely by a large portion of chromosome 9 (Duffin, 2024). DeBiasse et al. (2022) indicates a large inversion between the reference genome of *Pisaster brevispinus* and chromosome 9 of *P. ochraceus*; this inversion appears to be polymorphic in our data. Hereafter we exclude analysis of chromosome 9 and only focus on the remaining chromosomes.

For the PCA on the remaining neutral SNP dataset, the first PC axis explains ∼1.90% of the variation and separates individuals in the Salish Sea from those collected along the outer coast (Figure 2D). The second PC axis explained 1.72% of the variation and further separated individuals from DBW and FHW within the Salish Sea (Figure 2D).

**Figure 2.**
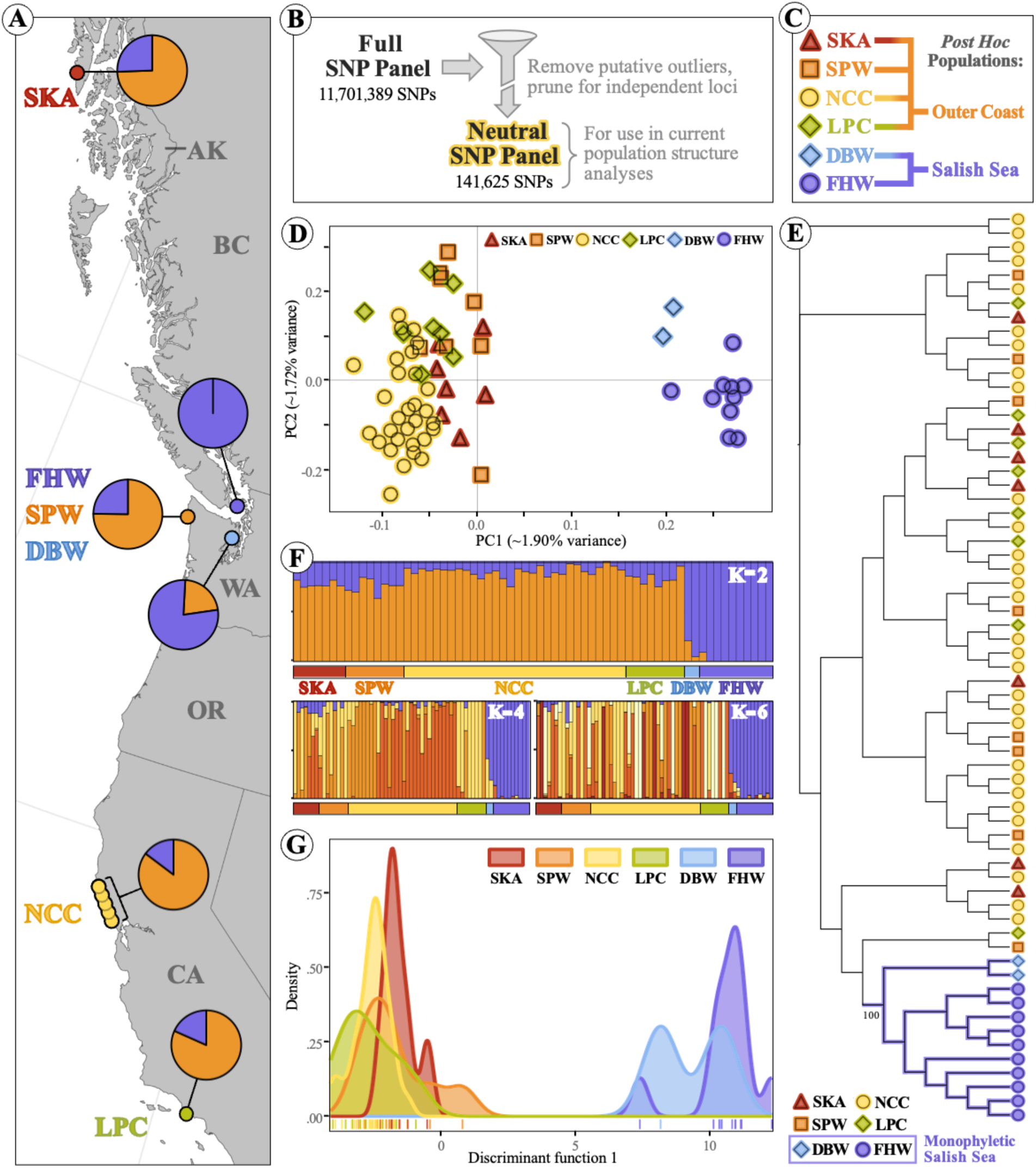
**Neutral population structure analyses including ADMIXTURE, PCA, sNMF, DAPC, and cladogram results**. Population structure analyses using the neutral panel (141,625 SNPs) generated from whole-genome ochre star SNP data. (**A**) Sampling map supplemented with ADMIXTURE results for K=2. Pie charts represent population membership across individuals at each sampling location. Full results for K=2 through K=10 run in triplicate and log likelihood score plot in supplement (Figure S5.1). (**B**) Overview of filtering pipeline used to create neutral SNP panel for current population structure analyses. (**C**) Color and shape legend for the six major sampling locations, applied consistently throughout all figures, which collapse into two *post hoc* groupings based on population structure results: outer coast and Salish Sea, represented by orange and purple shading, respectively. (**D**) Principal components analysis (PCA) scatterplot. PC1 values multiplied by −1 to match the geographical orientation of the outer Pacific coast relative to the Salish Sea. (**E**) Cladogram results displayed in FigTree. Bootstrap support value given as a percentage for the monophyletic Salish Sea clade, highlighted in purple, and calculated using Nei’s distance matrix, neighbor-joining tree. (**F**) sNMF (sparse Nonnegative Matrix Factorization) results for selected K values. Each x-axis bar represents one sea star’s membership to each of K populations, grouped by sampling location. Full results for K=2 through K=10 run in triplicate and cross entropy score plot in supplement (Figure S5.2). (**G**) DAPC (Discriminant Analysis of Principal Components) results, with x-axis tick marks represent DF1 (Discriminant Function 1) values per individual. DF1 values multiplied by −1 to again match geographical orientation.

Principal component analysis (PCA) of the OC-only full SNP panel did not reveal distinct clustering by sampling location, nor did it indicate substantial genetic differentiation among individuals regardless of origin (Supplement Figure S5.3). Individuals from different sites showed considerable overlap in PCA space, and genetic distances within and among sites were comparable, even between the most geographically distant locations (SKA and LPC) (Supplement Figure S5.3).

*Additional Population Structure Analyses: ADMIXTURE, sNMF, DAPC, and Cladogram* Multiple analyses of the neutral SNP dataset—ADMIXTURE (K = 2), sNMF (K = 2), DAPC, and a neighbor-joining phylogenetic tree based on Nei’s distance (Figure 2)—consistently revealed strong differentiation between individuals from the Salish Sea (DBW, FHW) and those from outer coast locations (SKA, SPW, NCC, LPC), mirroring the pattern observed in PCA. Further, sNMF-based population assignment among outer coast samples became increasingly variable and stochastic at higher K values (K = 3-5, 7-10; results not shown), whereas Salish Sea individuals consistently formed a distinct cluster across all values of K, underscoring a robust statistical separation. The absence of well-defined *post hoc* groupings among outer coast individuals indicates a lack of genomic structure by geography, ecotype, or other unmeasured factors.

#### No isolation-by-distance (IBD) among outer coast sites

Finally, we assessed isolation-by-distance (IBD) using a Mantel test among stars assigned to the OC population using the neutral SNP dataset, further separating the 5 subsampled regions of NCC into separate locations (NCC_1_ through NCC_5_, Supplement S6). The subsampled regions of NCC, all clustered within a ∼153 km region, spanned a larger range and overall magnitude of genetic distance than our two most disparate sites along the outer coast, SKA and LPC (separated by over ∼4170 km). This underscored a broader trend of no correlation between genetic and geographic (marine) distances (R^2^ = 0.00882, p = 0.38), suggesting no pattern of IBD among outer coast sites (Supplement S6).

##### Mitochondrial Genome Analysis

Mitochondrial nucleotide diversity was substantially lower than genome-wide diversity across all sampling locations (Supplement Table S4.1). Genome-wide π values ranged narrowly from 0.00850 to 0.00894, while mitochondrial values were markedly lower (0.00025-0.00211), representing ∼4- to ∼16-fold reductions in nucleotide diversity (excluding DBW, where the ∼35-fold estimate is likely inflated due to low sample size, n = 2; Table 1). *Post hoc* population groupings revealed the same pattern, with genome-wide π in the outer coast (0.00887) and Salish (0.00855) populations markedly higher than mitochondrial π in the outer coast (0.00178; ∼5x reduction) and, in particular, the Salish (0.00059; ∼14x reduction). As in previous studies (Harley et al., 2006; Marko et al., 2010) there is no evidence of spatial structure between the Salish samples and those on the outer coast (Supplement S7).

##### Outlier analyses identify putative selective sweep regions

To identify genomic regions potentially shaped by selection, we first applied a conventional outlier approach comparing pairwise F_ST_ with the log_2_ ratio of nucleotide diversity (π) between populations. This framework identifies regions that are both highly differentiated and asymmetrically reduced in diversity, pointing to candidate selective sweeps and the population in which diversity has been depleted (e.g., Chai et al., 2020; Li et al., 2020; Zhang et al., 2020).

Using this method on our full dataset (11,701,389 SNPs), we identified 65 genomic windows falling within the top 1% of F_ST_ values (≥ 0.107) and the top or bottom 1% of π ratio values (log_2_ π ratio ≥ 0.481 or ≤ –0.981) (Figure 3). Of these, 19 showed reduced diversity in the OC population and 46 in the SS population, indicating population-specific sweep signals.

**Figure 3.**
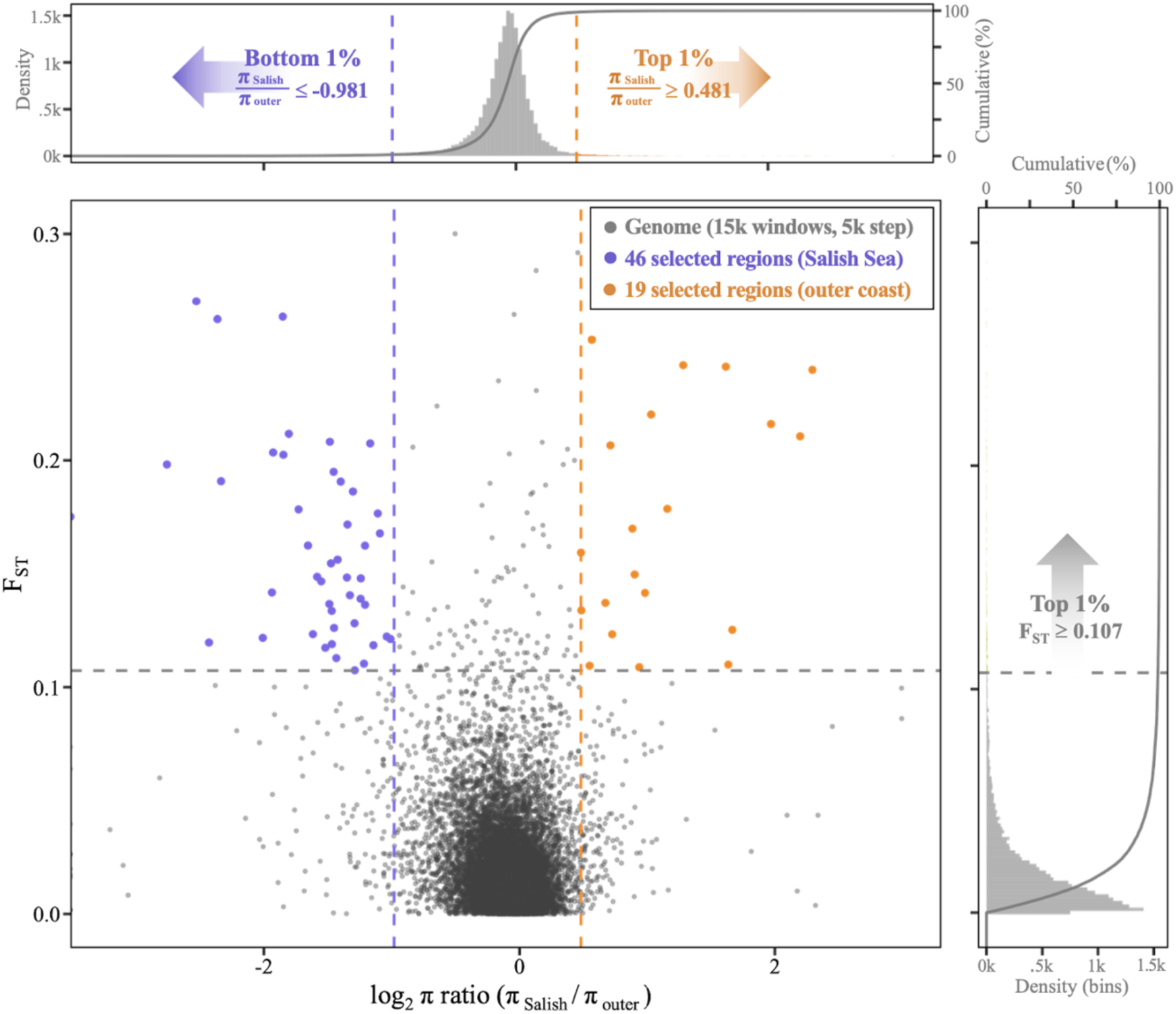
Genome-wide selective sweep plot comparing outer coast to Salish Sea individuals. Distributions of the π ratio (π _Salish_ / π _outer_, log_2_ transformed, displayed as histogram in top panel) and pairwise F_ST_ values (histogram distribution in right panel) were calculated in 15kb windows with 5 kb steps that were determined as optimal for the *P. ochraceus* genome size (see Table 2.3 for justification and literature review). The horizontal dashed gray line represents the threshold that selects for the top 1% of F_ST_ values, where F_ST_ ≥ 0.107. Vertical purple and orange dashed lines represent thresholds that select the bottom 1% and top 1% of log_2_ π ratios, where log_2_ (π _Salish_ / π _outer_) is ≤ −0.953 or ≥ 0.456, respectively. The top left and top right sectors formed by said threshold lines define candidate selective sweep regions (purple- or orange-shaded points) among Salish Sea or outer coast individuals, respectively.

To extend this framework, we developed a novel visualization approach, the “meadow” plot, which retains the genomic resolution of a Manhattan plot while incorporating the directionality provided by π ratio. By plotting F_ST_ values by chromosomal position and coloring windows based on π ratio, meadow plots allow rapid identification of outlier regions and the population likely driving the signal. When applied to all pairwise site comparisons and a direct comparison of OC and SS populations (Figure 4; Supplement S8), meadow plots revealed distinct clusters of candidate sweep regions, highlighting divergent evolutionary pressures across the range.

**Figure 4.**
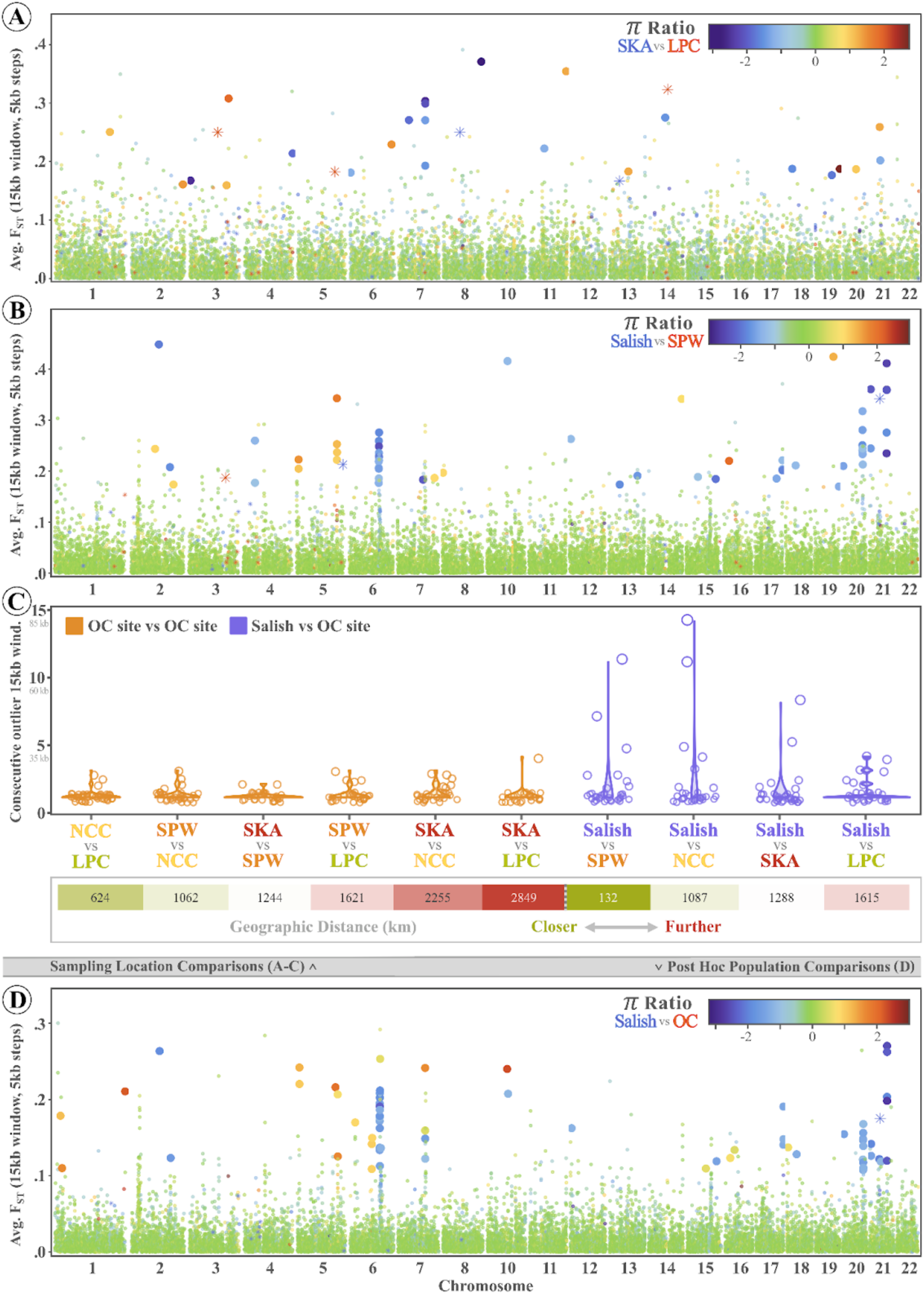
Selected meadow plot results of interest displayed as genomic regions of outlier F_ST_ and log_2_ π ratios across 15kb windows (with 5kb steps). (**A**) Selected sampling location-based meadow plot comparing OC sites SKA and LPC and representing the two most geographically-distant sampling locations at 2849 km apart. (**B**) Selected sampling location-based meadow plot comparing OC site SPW and Salish Sea-collected individuals (following FHW and DBW sample merge) to represent the stark divide between OC and Salish individuals, despite relative geographic proximity (132 km). (**C**) Violin plots summarizing the spread of consecutive outlier windows for each of the 10 pairwise sampling locations. Orange-colored violin plots denote instances where one OC site is being compared to another, while purple-colored violin plots represent an OC site being compared to the Salish Sea (i.e., FHW and DBW combined). (**D**) Meadow plot comparing post hoc populations defined through population structure analyses, i.e., the Salish Sea population (FHW and DBW) compared to the outer coast population (SKA, SPW, NCC, and LPC). Across all meadow plots, windows falling within both the top 1% of FST and the top or bottom 1% of log_2_ π ratios are considered outliers (defined using the same criteria pattern as described in Figure 3 caption), with datapoints representing outlier windows scaled up in size for emphasis. Finally, asterisk-shaped datapoints represent comparisons where one of the two populations under comparison had a π value of zero, resulting in a mathematical error calculation (+/-inf) for the log_2_ π ratio.

Some pairwise comparisons, such as SKA–LPC, showed a relatively balanced mix of SKA- and LPC-driven outliers across the genome (Figure 4A). Others, including Salish–SPW, were dominated by cool-toned signals, suggesting that most sweep regions occurred in Salish-collected individuals (Figure 4B). This Salish-driven pattern was consistent across all four Salish–OC comparisons, with recurrent peaks corresponding to large spans of consecutive outlier windows on chromosomes 6 and 21 (Supplement S8). Indeed, the *post hoc* population based meadow plot reflects the same trends displayed individually among each of the four Salish-OC contrasts (Figure 4D).

To quantify peak strength, we defined intensity as the number of consecutive windows that qualified as both F_ST_ and π-ratio outliers per region. This revealed that the strongest sweep signal (14 consecutive outlier windows) occurred in the Salish vs. SPW comparison (Figure 4C). In general, Salish–OC comparisons showed more and stronger peaks than OC–OC comparisons. Because outliers were defined separately for each pairwise comparison, these results likely underestimate the true extent of Salish–OC divergence, as a universal threshold would highlight even fewer OC–OC outliers.

## Discussion

The limited spatial diversity amid outer coast populations of *P. ochraceus* reaffirms previous work (Harley et al., 2006, Marko et al., 2010, Schiebelhut et al., 2022a) and offers a spatial contrast to the apparent isolation of the Salish locales in our study. As a free-spawning species with planktotrophic larvae and an extended pelagic larval duration (Strathmann, 1978; Pia et al., 2012), *P. ochraceus* possesses traits that facilitate dispersal and should theoretically translate into high gene flow. Additionally, the perceived boom-bust demographic cycles of asteroids (Uthicke et al., 2009) may further limit spatial genetic structure through recolonization dynamics (Schiebelhut et al., 2022b). Indeed, earlier genetic studies investigating population structure in *P. ochraceus* supported this expectation, finding evidence of extensive gene flow across major latitudinal and ecological gradients (Harley et al., 2006; Schiebelhut et al., 2022a). Our findings show for the first time pronounced spatial population structure in *P. ochraceus* when comparing Pacific open-coast populations to those within the Salish Sea. Based on whole-genome data with only putatively neutral sites included, there is statistically robust isolation between these regions; with all data included, distinct contiguous shifts in genomic diversity emerge, suggesting a potential role for local adaptation in shaping this divergence.

Evolution does not act uniformly across the genome; while some regions are rapidly shaped by strong selective pressures, many others remain effectively invisible to selection, evolving more gradually as functionally silent mutations arise and are passively carried by genetic drift (Kimura, 1979). Variation in historical demography, including range expansions, can further shape the diversity of a species across its distribution (Marko & Hart, 2011; Dawson, 2013).

These combined forces may lead to marine species exhibiting population structure that may deviate from our predictions based on dispersal potential or other life history traits alone (Zakas & Wares, 2012; Dawson et al., 2014; Eldon et al., 2016; Guo & Wares, 2017). This complexity makes it essential to distinguish adaptive variation from patterns driven by historical and ongoing barriers to gene flow (e.g., Xuereb et al., 2021). We first analyzed population structure using a putatively neutral SNP panel to minimize selection signals and provide a clearer view of demographic history and connectivity in *P. ochraceus*. This framework then informed our downstream analyses of adaptive differentiation, ensuring that signals of selection were interpreted within the broader context of population structure and gene flow dynamics.

While the environmental differences between the Pacific coastal ecoregions and the Salish Sea are substantial and likely introduce selective pressures, the patterns we present suggest a combined influence of geographic isolation and local adaptation. Although selection can indirectly affect the frequencies of neutral loci linked to strongly selected variants through genetic hitchhiking, the genomic signatures of this process appear as a limited number of distinct, localized selective sweeps, each confined to a narrow portion of a chromosome. In contrast, the unlinked neutral loci contributing to our signal of spatial isolation are broadly distributed throughout the genome, suggesting that some degree of dispersal limitation has shaped genetic differentiation between Salish stars and those along the outer coast, even if the divergence remains relatively recent on a geological timescale (explored below). Taken together, the persistence of a genome-wide divergence signal in loci unaffected by selection underscores the power of whole-genome data to detect subtle or recent constraints on gene flow. Further, we emphasize the value in distinguishing between neutral and adaptive variation in genomic analyses to uncover subtle signatures of restricted gene flow and the influence of both historical and ongoing dispersal barriers.

We developed a novel approach to highlight putative selective sweep regions across the *P. ochraceus* genome by integrating classic Manhattan plots with selective sweep detection analyses. The latter is an increasingly utilized approach (e.g., Chai et al., 2020; Li et al., 2020; Zhang et al., 2020) that identifies candidate selective sweep regions by plotting pairwise genetic differentiation (*e.g.*, F_ST_) against the ratio of nucleotide diversity (π) between populations. The π-ratio statistic offers directionality by indicating which population shows unexpectedly low diversity, revealing likely drivers of sweep signatures. Two key observations emerge from this approach. First, distinct chromosomal regions exhibit strong evidence of selection across multiple pairwise population comparisons, with trends primarily pointing to the Salish Sea as exhibiting reduced nucleotide diversity signals underlying outlier behavior. Second, these regions frequently span >100 kb, suggesting recent, strong selection episodes in the Salish population, a pattern indicative of adaptation to novel environmental pressures (Villegas-Mirón et al., 2021; Rogers et al., 2023).

The prevalence of large-scale sweep regions in the Salish Sea suggests that *P. ochraceus* populations in this semi-enclosed estuarine system have undergone relatively recent adaptation to its distinct abiotic conditions. Unlike the outer Pacific coast, the Salish Sea is characterized by strong seasonal fluctuations in salinity, increased freshwater input, and differences in nutrient and trace metal availability, particularly dissolved iron levels, all of which can exert significant selective pressures on marine organisms (Khangaonkar et al., 2019; Silliman, 2019; Gierke et al., 2023). Adaptive divergence in response to such conditions has been observed in other taxa, such as *Gadus morhua* populations in low-salinity fjords (Berg et al., 2015) and *Homarus americanus*, where ocean currents and thermal gradients structure adaptive genetic variation (Benestan et al., 2016). Some studies even show phenotypic shifts in feeding behavior of *P. ochraceus* related to the difference in salinity between these two regions (Held & Harley, 2009; Pia et al., 2012).

While neutral genetic structure indicates some degree of dispersal limitation, the extensive selective sweeps suggest that *P. ochraceus* populations within the Salish Sea have further been shaped by selection pressures distinct from those acting on outer coast populations.

Given the strong environmental differences across sites, we conducted preliminary redundancy analyses (RDA) to explore potential genotype-environment associations (see Supplement S9). However, because we lack replication across multiple demographic demes that span similar environmental gradients, it’s difficult to determine whether the observed genetic differences reflect environmental selection or simply isolation. As such, while our findings strongly suggest a role for environmental selection, additional studies incorporating denser sampling across transitional zones, common-garden experiments, or genome-environment association approaches with stronger replication are needed to disentangle selection from potential confounding demographic effects.

The value of the meadow plots in Figure 4 lies in their ability to narrow the set of target regions for follow-up; regions with elevated F_ST_ but minimal differences in diversity can be prioritized differently from those showing elevated F_ST_ alongside a pronounced reduction in diversity in one population. To illustrate the utility of our meadow plot approach, we highlight a pronounced region of divergence on chromosome 6, spanning ∼180 kb, where nucleotide diversity is markedly lower in the Salish Sea relative to the outer coast. Examination of this ∼180kb “peak” region via the top BLAST hits highlights orthologs of cytochrome P450 in *Asterias* (LOC117288746) and *Patiria* (LOC119736944) as well as histone methyltransferase/deacetylase annotations (LOC117295344, LOC117306839). Both protein types are reliant on dissolved iron as a cofactor (Sharifian et al., 2020, Shapiro et al., 2023), and echinoderms are thought to be sensitive to environmental metal concentrations, particularly iron (Gonzáles-Delgado et al., 2024). Given the iron-rich runoff from the rivers leading into the Salish Sea (Williams & Chan, 1966; Chase et al., 2007) and the strength of signature for iron as a driver in our RDA analysis (Supplement 9), this may be an intriguing candidate region for more focused work on mechanisms of adaptation between the coastal and Salish populations of *P. ochraceus*.

This apparent population break has implications for *P. ochraceus* as climate change progresses; warming in Pacific coastal waters is expected to outpace that of the Salish Sea (Khangaonkar et al., 2019). The fitness landscape for *P. ochraceus* across a range of temperatures has only been locally evaluated in the Salish Sea (Vickery & McClintock, 2000; Gooding et al., 2009; Walton et al., 2024), with no comparable studies on the outer coast. One way or the other, the same isolating mechanism(s) limiting gene flow today may prevent open-coast stars from successfully colonizing the Salish Sea, with potential fitness costs if dispersal does occur. Both the underlying cause of these barriers and their impact on future ochre star persistence remain open questions, yet they may reflect wider processes affecting marine species of the region.

The observed divergence between *P. ochraceus* populations in the Salish Sea and the outer Pacific coast is likely shaped by a combination of oceanographic, environmental, and historical factors. Taken together, our genomic evidence suggests that a founder effect may have influenced the genetic diversity of the Salish Sea population, supporting the hypothesis that this region was colonized relatively recently by migrants from the outer coast following deglaciation of the region less than 13,000 years ago (Clague, 1989; Buonaccorsi et al., 2002). Further, while several uncertainties complicate the timeline estimation of Salish stars’ demographic history, the net nucleotide divergence between samples from Friday Harbor and the outer Washington coast is low (∼0.00008). Given current mutation rate estimates from other asteroids (Popovic et al., 2024) coupled with the patterns we observed across neutral and adaptive SNPs, we posit that a post-glacial founder event may have seeded this divergence. Infrequent and/or episodic gene flow has likely further obscured the timing of this event, with mitochondrial genomes appearing notably homogeneous across the range of *P. ochraceus* (Harley et al., 2006; Supplement S7); discordance between nuclear and mitochondrial patterns is not uncommon (Toews & Brelsford, 2012).

Regardless of the timing of *P. ochraceus*’ arrival in the Salish Sea, genomic evidence clearly implicates local adaptation coupled with some degree of physical isolation as key drivers of the observed divergence that exceeds what would be expected from geographic distance alone. The Strait of Juan de Fuca, a glacially formed fjord, serves as the primary conduit for exchange between the river-fed Salish Sea and the marine waters of the Pacific, yet strong estuarine outflow creates a persistent salinity gradient and limits passive larval transport into the interior (Herlinveaux & Tully, 1961; Khangaonkar et al., 2019; see Wares, 2025). Moreover, the Salish Sea itself is highly structured, consisting of two main basins (the Strait of Georgia and Puget Sound) along with a complex network of smaller basins and fjords. These basins are delineated by bathymetric sills (e.g., Ebbesmeyer et al., 1984; Sobocinski, 2021), which contribute to long water residence times and, in some taxa, population substructure within the region (e.g., Dungeness crab, Jackson & O’Malley, 2017; Greenstriped Rockfish, Wray et al., 2025). Of particular relevance to exchange between the outer Pacific and the Salish interior is the Victoria Sill, a shallow (55–100 m) submarine ridge spanning the inner portion of the Juan de Fuca Strait (Ebbesmeyer & Barnes, 1980; Khangaonkar et al., 2017; Wray et al., 2025). This hydrographic feature has been shown to influence circulation and water retention (e.g., Ebbesmeyer et al., 1984), potentially restricting larval influx from the outer coast while further structuring salinity gradients that may drive selection in resident populations (e.g., Masson & Cummins, 2000; Lawlor & Arellano, 2020).

Our *a priori* sampling design, guided by previous studies, was not optimized to delineate the geographic boundary of this genomic transition. Instead, our resolution is limited to a ∼200 km stretch between the San Juan Islands (Friday Harbor, WA; FHW in Figure 1) and the outer Olympic Peninsula (Chilean Memorial at Sokol Point, WA; SPW in Figure 1). Finer-scale sampling, particularly throughout the Juan de Fuca and across the Victoria Sill, is needed to clarify the role of dispersal limitation in shaping these patterns. Community-level patterns offer additional insight as similar reports of spatial divergence into the Salish Sea are becoming increasingly common (e.g., Wray et al., 2025), suggesting that the genetic transition we observe in *P. ochraceus* is not an isolated phenomenon but part of a notable ‘microbiogeographic’ feature. Many species exhibit spatially concordant genetic discontinuities across the same geographic region, pointing to shared barriers that shape population structure across taxa (Box 1).

Unlike the strong genetic break between the Salish Sea and the outer Pacific, *P. ochraceus* populations along the outer coast from southern Alaska to southern California show no statistical evidence of spatial structure, including patterns expected under isolation by distance, indicating high connectivity across more than 2,800 km of coastline. The strong alongshore transport facilitated by the California Current System and regional upwelling undoubtedly enhances connectivity among outer coast sites (Pringle & Wares, 2007; Wares & Pringle, 2008) and likely inhibits detectable divergence at neutral loci (Palumbi, 2004; Cowen & Sponaugle, 2009).

Specifically, the large population size of high-dispersal outer coast stars, combined with their propensity for demographic fluctuations (Uthicke et al., 2009) and metapopulation structure (Schiebelhut et al., 2022), weakens the impact of genetic drift in driving differentiation. In this context, even low-level or sporadic gene flow acts to spatially homogenize sequences, gradually eroding signals of divergence (Andrews, 2010) or leading to an appearance of ‘chaotic genetic patchiness,’ more accurately described as ‘eurymixis’ (Dawson et al., 2011).

Failure to resolve fine-scale population structure among outer coast sites in our neutral SNP panel does not preclude the possibility that adaptive or even demographic differences exist. While divergence is unpronounced among selectively neutral sites, there are clear regions of likely adaptive significance among these Pacific coast populations (Figure 3,4; Supplement S8).

Many marine taxa with high dispersal potential exhibit subtle but functionally important divergence that may be driven by environmental gradients rather than geographic isolation (Sanford & Kelly, 2011; Zerebecki et al., 2021; Sotka et al., 2024), and the patterns of trait adaptation may vary broadly throughout a species range (Hice et al., 2012). The specific role of particular genomic regions in adaptive divergence is still difficult to assess in non-model species such as *P. ochraceus*, as gene annotations are indirectly applied from experimental organisms that are evolutionarily distant. Future work incorporating larval dispersal modeling or environmental association analyses may help determine whether localized selection is shaping population-level variation despite high gene flow along the exposed Pacific coastline. For now, these results add context to decades of work on life history and ecological variation across the distribution of one of the best-studied and charismatic marine predators.

## Acknowledgements

Bob Schmitz offered phenomenal assistance with Illumina library handling at UGA. Thanks to Jeff Ross-Ibarra for numerous great ideas throughout this project and Erik Sotka for initial comments on the manuscript. Shan-Ho Tsai at Georgia Computational Resource Center deserves a big raise. Work supported by the National Science Foundation (OCE-1737091) to JPW and (OCE-1737381) to MND, with gratitude to our NSF program officers and colleagues for tremendous service to the scientific effort of our nation. We thank Sabrina Pankey, Peter Raimondi, Melissa Miner, Maya George, CJ Urnes, Maya George, Melissa Douglas, Shawn Knapp, and Jesse Wilson for their help collecting tissue samples.

## Funding

This work was supported by NSF-OCE-1737091 to JPW and NSF-OCE-1737381to MND.

## Data Availability

Sequence data are at NCBI SRA under BioProject 1117092 and NSF BCO-DMO datasets at DOI: 10.26008/1912/bco-dmo.934772.1 (Wares and Duffin 2024). Code is at <u>github.com/paigeduffin/PhD.work_WGS.chapter</u>

## Author Contributions

PJD, MND, LMS, and JPW designed and performed the research, analyzed the data, and wrote the paper.

### Box 1

**Multi-species analysis of divergence across the Salish-Pacific interface.**

The Salish Sea functions as a vast and dynamic estuary (Sobocinski, 2021), where tidal inflow of marine water from the Pacific mixes with freshwater outflow (Herlinveaux & Tully, 1961), suggesting that populations of marine organisms could be well connected across this interface. Nevertheless, the endemicity of biodiversity in the Salish is striking, with divergent populations or species across multiple taxa, including marine mammals (Barrett-Lennard & Ellis, 2001; Morin et al., 2024), macroalgae (Gierke et al., 2023), and phytoplankton (Whittaker & Rynearson, 2017).

The unique configuration of the Salish Sea, which has been profoundly shaped by glaciation (Thorson, 1980; Booth et al., 2003), creates stark contrasts in macroinvertebrate diversity relative to the outer Pacific coast. In the figure at right, we illustrate data from a review of population connectivity studies for marine species with pelagic larvae with sampling ranges spanning this region. Of 19 species meeting these criteria, 11 show support for genetic isolation in the Salish Sea (supported by K-means methods, e.g., STRUCTURE, and/or elevated divergence metrics relative to distance) and cover a wide range of phylogenetic diversity (see Supplement S10 for details and references).

For many marine species, realized dispersal is influenced by a combination of intrinsic traits – Studies to date exhibiting clear support for genomically distinct populations between Salish Sea and outer coast Pacific. Inset: Pelagic larval duration (PLD) estimates from literature alone do not explain distinct patterns of apparent connectivity across taxa. Colors distinguish species and corresponding taxonomic groups. Map from SimpleMappr (Shorthouse, 2010). Data and references for plot in Supplement S10. reproductive timing (Byers & Pringle, 2006), larval behavior, and adult movement – and environmental or physical barriers that thus may have species-specific effects on connectivity. Understanding these factors is key for interpreting the evolutionary ecology of organisms with dispersal that is primarily during the larval phase. Pelagic larval duration (PLD) is often considered a predictor of dispersal capacity and genetic structure. However, many species illustrate substantial population structure across this region despite extended PLD, as noted in Wray et al. (2025). Even a species with modest PLD (∼10 days, *C. nuttalli*), well above the typical threshold for ‘extended’ PLD (Shanks, 2009; Pappalardo et al., 2015), exhibits this divergence. Tracking larval traits other than PLD, e.g. diel migration or salinity effects on feeding, remains challenging due to their context dependency (Strathmann, 1978; Pineda et al., 2007; Pia et al., 2012; Hodin et al., 2019). Additionally, the mechanisms driving population diversity and divergence across a ‘seascape’ are notably complex (Marko & Hart, 2011; Dawson, 2013). The species that do not show notable divergence across this boundary may have a more recent demographic expansion into the Salish Sea region.

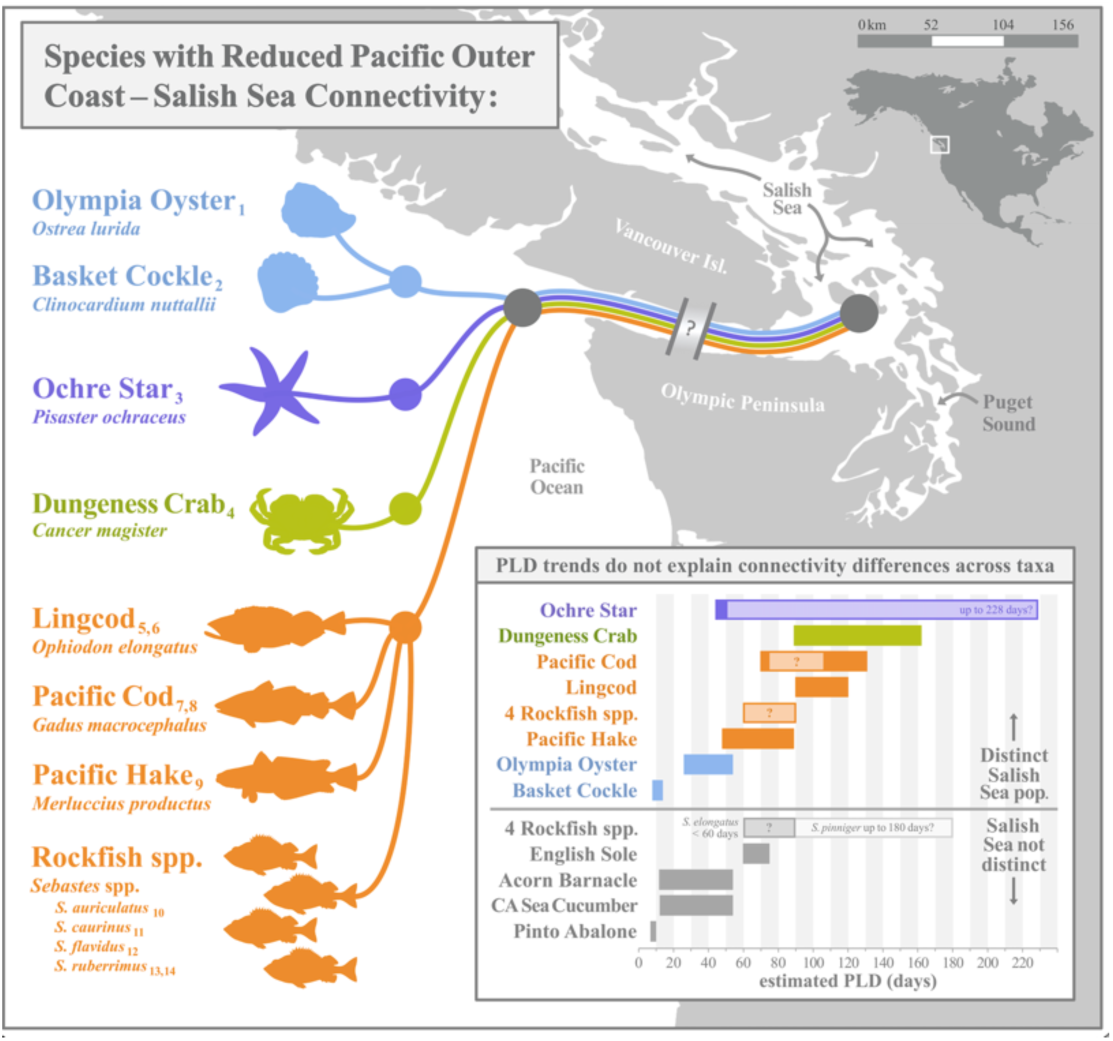

## Supplemental Information for

### Supplement S1. Latitudinal temperature gradient

**Figure S1.1.**
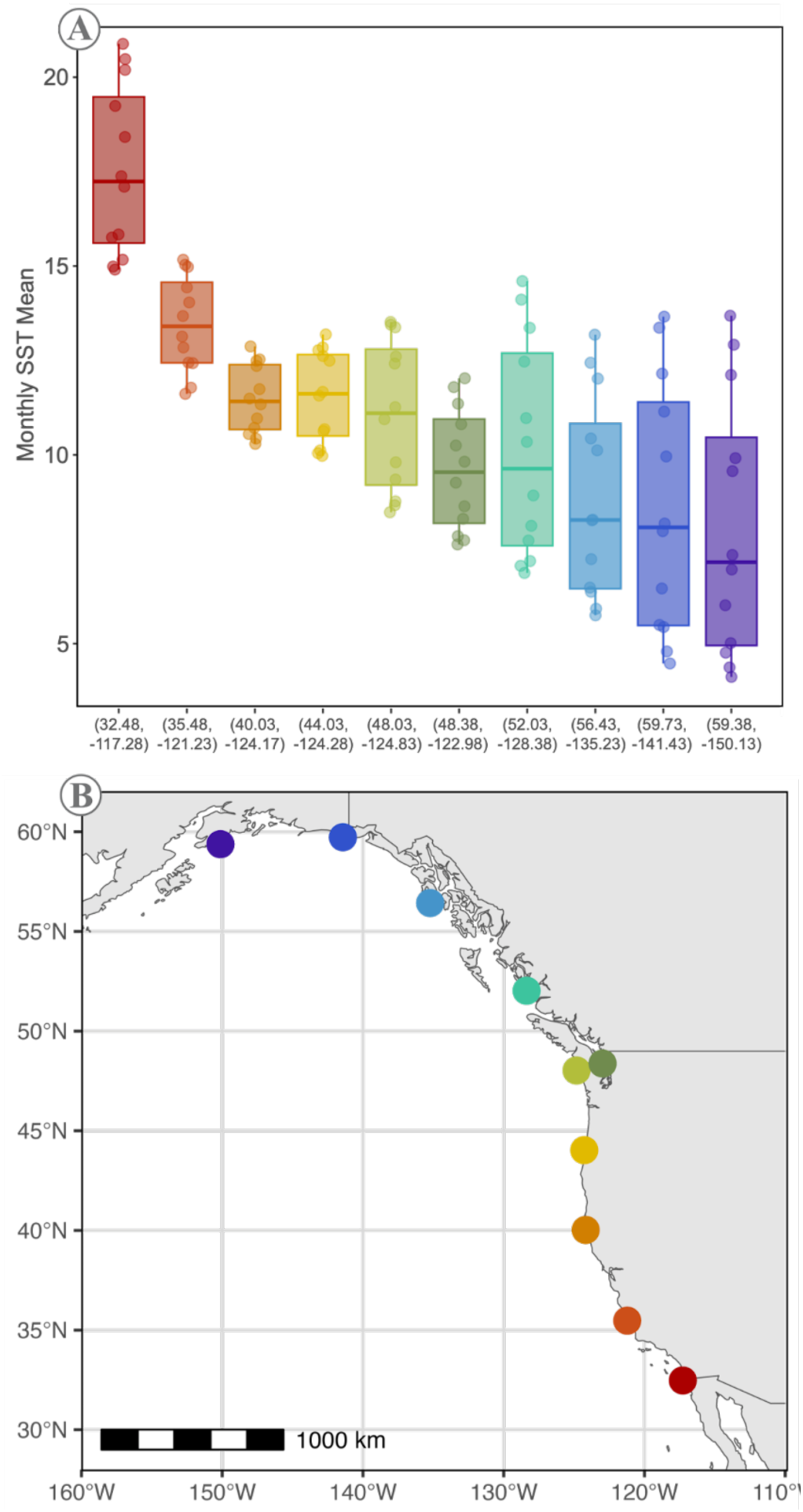
(A) Monthly sea surface temperature (SST) mean values in 2013 at (B) nine offshore and one inshore (Salish Sea) sampling locations of interest. along the west coast of North America encompassing the approximate species range of *Pisaster ochraceus*.

#### Methods

Monthly sea surface temperature (SST) mean values were extracted as in Dawson et al. (2023) from the “SST and SST Anomaly, NOAA Global Coral Bleaching Monitoring, 5km, V.3.1,

Monthly, 1985–Present” dataset at National Oceanic and Atmospheric Administration Environmental Research Division Data Access Program (NOAA ERDDAP; https://coastwatch.pfeg.noaa.gov/erddap/griddap/NOAA_DHW_monthly.html), then filtered to represent one year of interest (2013). Then, monthly SST mean values were plotted at nine approximately equidistant offshore and one inshore (Salish Sea) sampling locations of interest using RStudio (RStudio Team, 2020).

## Supplement S2. Bioinformatics details

### Sequence Processing / SNP Calling Details

Paired-end reads were trimmed using Trim Galore! (Krueger, 2015) using the paired-read setting (trim_galore --paired {readA} {readB}). FastQC reports (Andrews, 2010; module version: FastQC/0.11.9-Java-11) were generated for each trimmed read file (fastqc {read}). The *P. ochraceus* reference genome (Hu et al., *in pre*) was obtained from collaborators and indexed with BWA (Li & Durbin, 2009; module version: BWA/0.7.17-GCC-8.3.0; command: bwa index p.ochraceus_genome.fna.gz). Paired reads were mapped to the reference genome using BWA MEM and stored as a .sam file (bwa mem -t 8 p.ochraceus_genome.fna.gz {readA}.fq.gz {readB}.fq.gz > {sample_ID}.sam). Next, sam (and later, bam) files were processed through a series of steps recommended in the Genome Analysis Toolkit (GATK) workflow (Broad Institute, DePristo et al., 2011): (**1**) duplicates were removed using the Picard (https://broadinstitute.github.io/picard/) command AddOrReplaceReadGroups; (**2**) sequences within each file were sorted with samtools sort (Li et al., 2009; module version: SAMtools/0.1.19-foss-2019b); (**3**) paired-end information was verified/corrected with Picard’s FixMateInformation command and output as bam files; finally, (**4**) duplicate reads were marked with Picard’s MarkDuplicates command.

The processed bam files were then used to call variants with a series of BCFtools (Li et al., 2009; module version: BCFtools/1.6-foss-2019b) commands (bcftools mpileup -Ou -f p.ochraceus_genome.fna --bam-list list.txt | bcftools call -mv -Oz -o vcf_file.vcf.gz); the resulting vcf (variant call format) file was indexed (tabix -p vcf vcf_file.vcf.gz) and used to generate a consensus sequence (bcftools consensus -f p.ochraceus_genome.fna vcf_file.vcf.gz > consensus.fa). Finally, VCFtools’ vcf-stats command (Danecek et al., 2011; version number: 0.1.16-GCC-8.3.0-Perl-5.30.0) was used to characterize basic statistics of the data prior to quality filtering.

**Figure S2.1.**
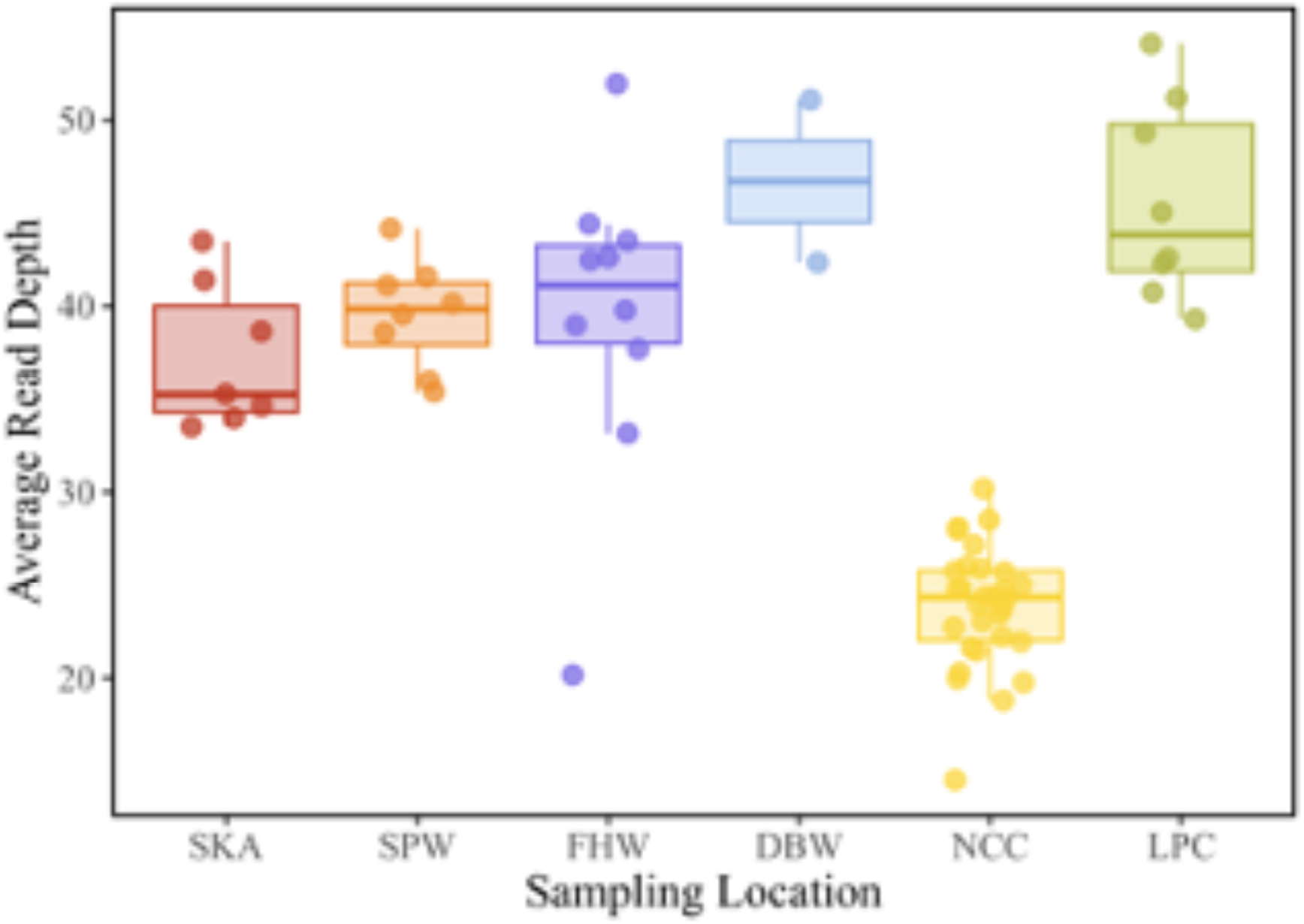
Average read depth by major sampling location. Each point represents the average read depth of one *P. ochraceus* individual sampled within that location. The lower average read depth among NCC-collected individuals reflects divergent sequencing pipelines. See methods for additional details.

**Table S2.1.** Methods used to quality-filter SNPs used in this analysis.

| Quality Filtering Step | SNPs Retained | SNPs Removed | Percent Removed |
| --- | --- | --- | --- |
| 1. [Variants, pre-filtering] | 26,340,914 | ---- | ---- |
| 2. Remove insertions/deletions | 26,340,914 | ---- | ---- |
| 3. Keep SNPs w/ < 20% missing across stars | 25,293,613 | 1,047,301 | 3.98% |
| 4. Keep SNPs w/ QUAL > 200 | 20,689,701 | 4,603,912 | 18.20% |
| 5. Keep SNPs w/ for 200 < DP < 2500 | 19,338,430 | 1,351,271 | 6.53% |
| 6. Keep SNPs w/ MQ > 40 | 18,337,510 | 1,000,920 | 5.18% |
| 7. Keep SNPs w/ < 10% missing across stars | 17,862,933 | 474,577 | 2.59% |
| 8. Keep SNPs w/ MAF > 5% | 11,701,389 | 6,161,544 | 34.49% |
| **Overall** | **11,701,389** | **14,639,525** | **55.58%** |

**Table S2.2.**
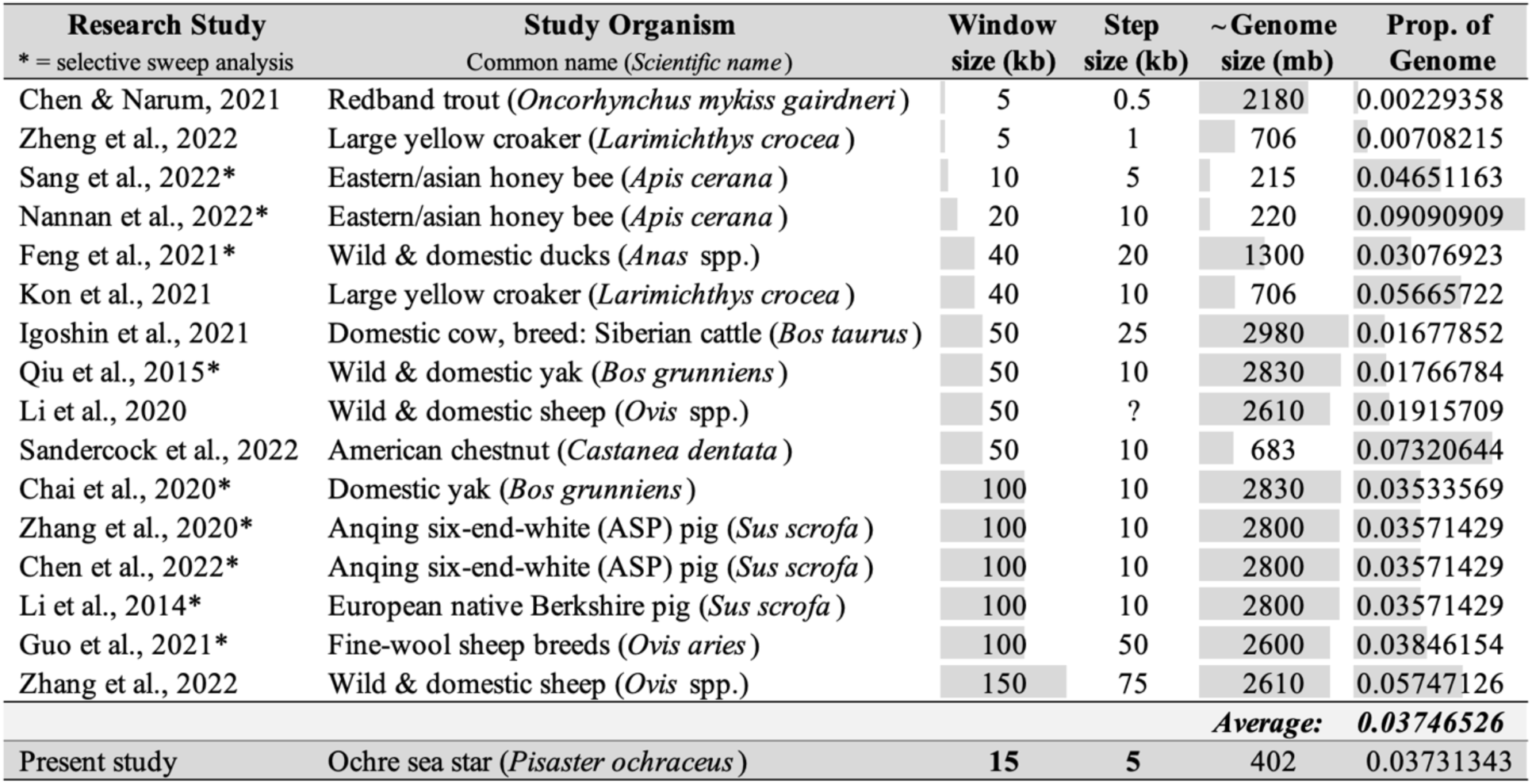
Justification for 15kb window bins used in this analysis. Note: genome size represents a time-efficient search and should not be used as a reference beyond this context, as some may, instead, represent a similar species whose information was more readily available at the time.

---

**Figure S2.2.**
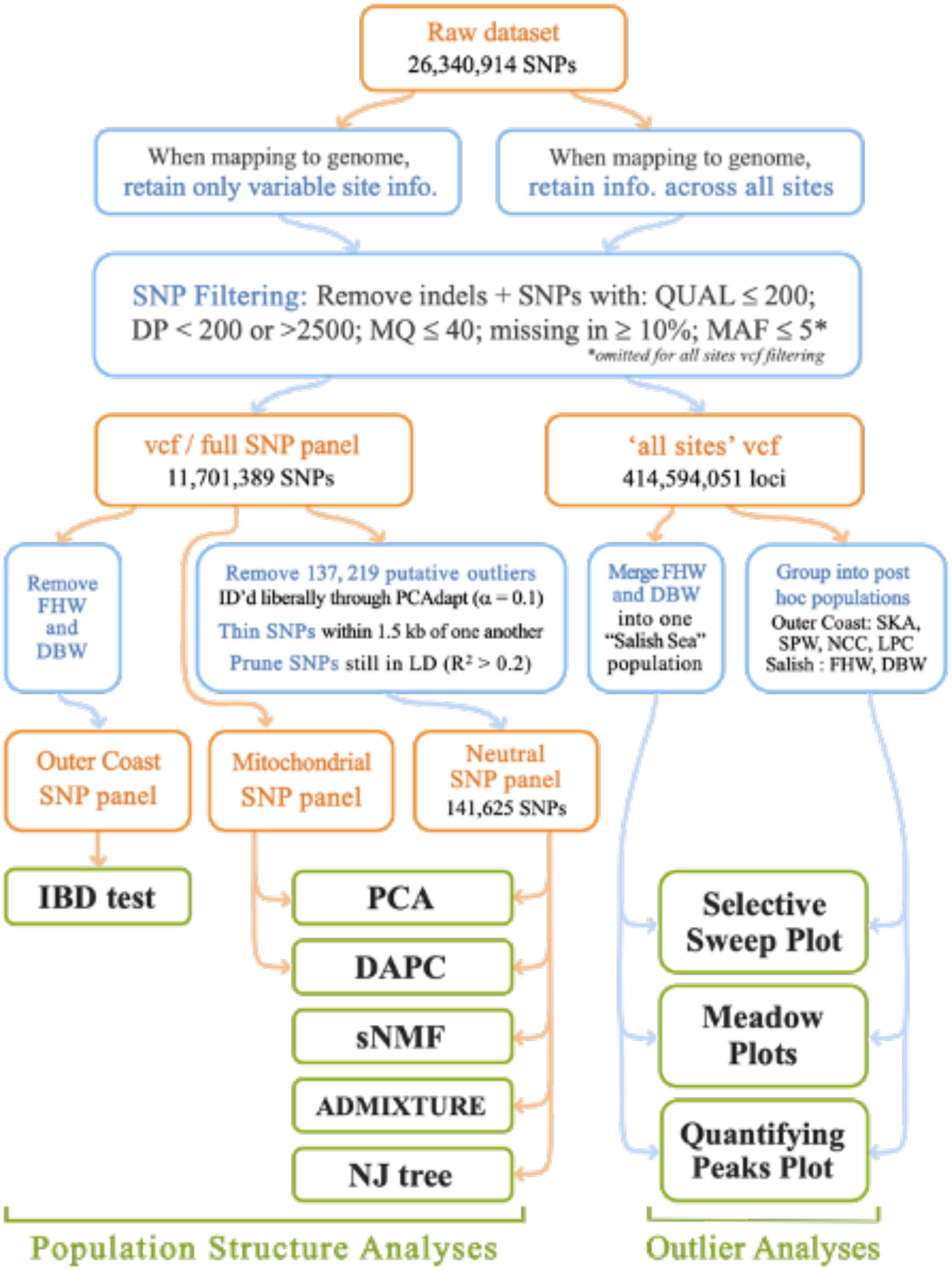
Variant filtering pipeline and subsequent analyses overview. Orange colored lines/font represents datasets and/or SNP panels, while filtering steps and analyses are in blue and green, respectively.

## Supplement S3. Linkage Disequilibrium

**Figure S3.1.**
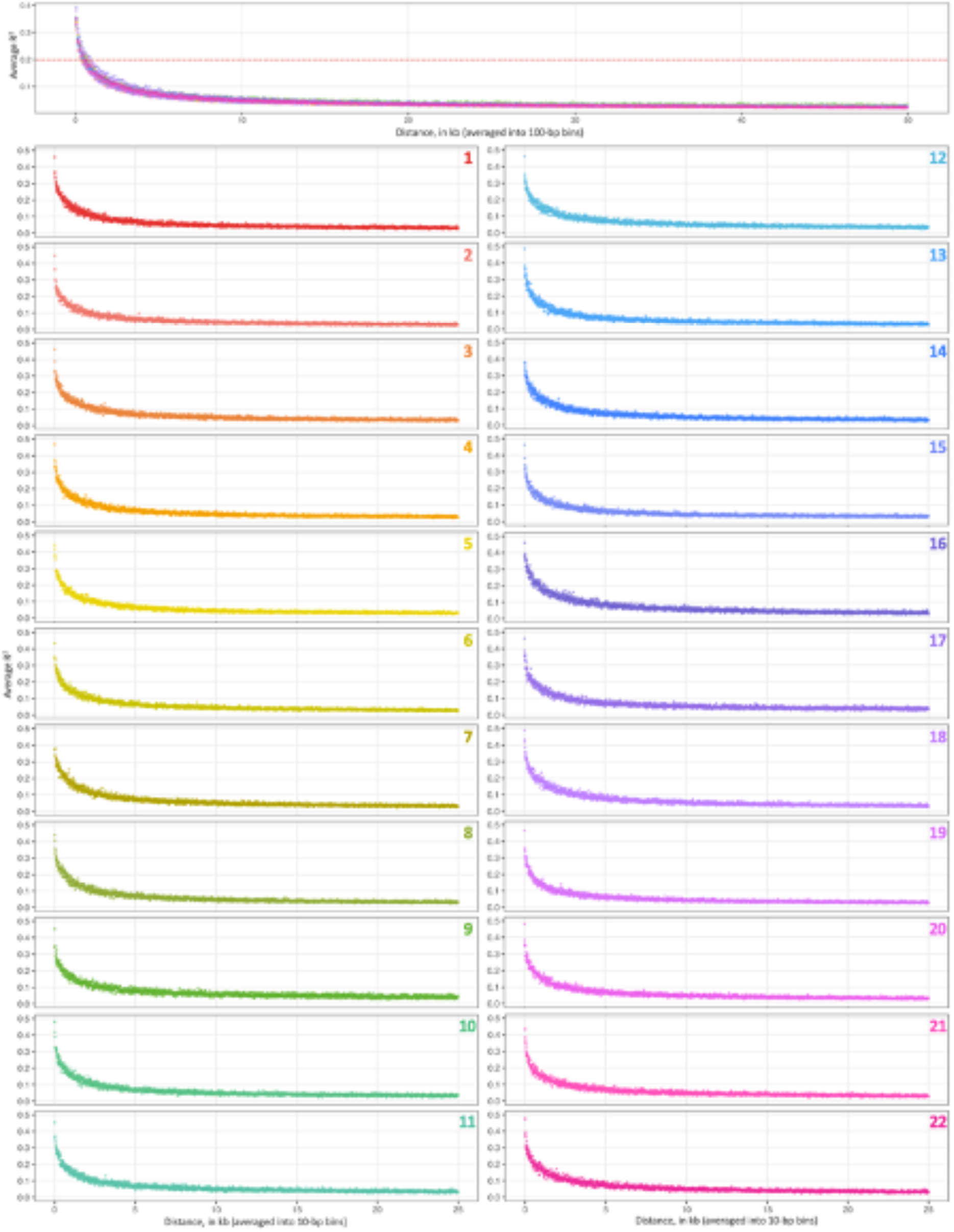
Linkage disequilibrium (LD) across all chromosomes (top panel) and by chromosome (panels 2 through 23) averaged across 10-bp bins (x-axis) and represented as the average squared correlation (R^2^) between a pair of loci (y-axis). Red dashed line in the top panel represents the LD value chosen as the cutoff for neutral SNP filtering (i.e., R^2^ = 0.2).

### Methods

We calculated linkage disequilibrium separately for each of the main 22 chromosomes using VCFtools (v 0.1.16) (Danecek et al., 2011) and the following generalized code: vcftools --gzvcf quality.filt_SNPs.vcf.gz --geno-r2 --ld-window-bp 10 --out LD_calc_10.bp.windows.

## Supplement S4. Genome-wide π diversity, variation

**Table S4.1.** A contrast of genome-wide π at each sample location relative to mitochondrial π. calculated for each major sampling location (rows 1-6) and post hoc population clusters ‘Salish’ (Salish Sea; FHW and DBW-collected samples) and ‘Outer’ (outer Pacific coast; SKA, SPW, NCC, LPC samples).

| Pop. | Genome-wide | Mitochon. | Magnitude Difference |
| --- | --- | --- | --- |
| SKA | 0.00890 | 0.00211 | ~ 4x |
| SPW | 0.00886 | 0.00057 | ~ 16x |
| NCC | 0.00883 | 0.00211 | ~ 4x |
| LPC | 0.00894 | 0.00140 | ~ 6x |
| DBW | 0.00863 | 0.00025 | ~ 35x |
| FHW | 0.00850 | 0.00061 | ~ 14x |
| Outer | 0.00887 | 0.00178 | ~ 5x |
| Salish | 0.00855 | 0.00059 | ~ 14x |

**Table S4.2.**
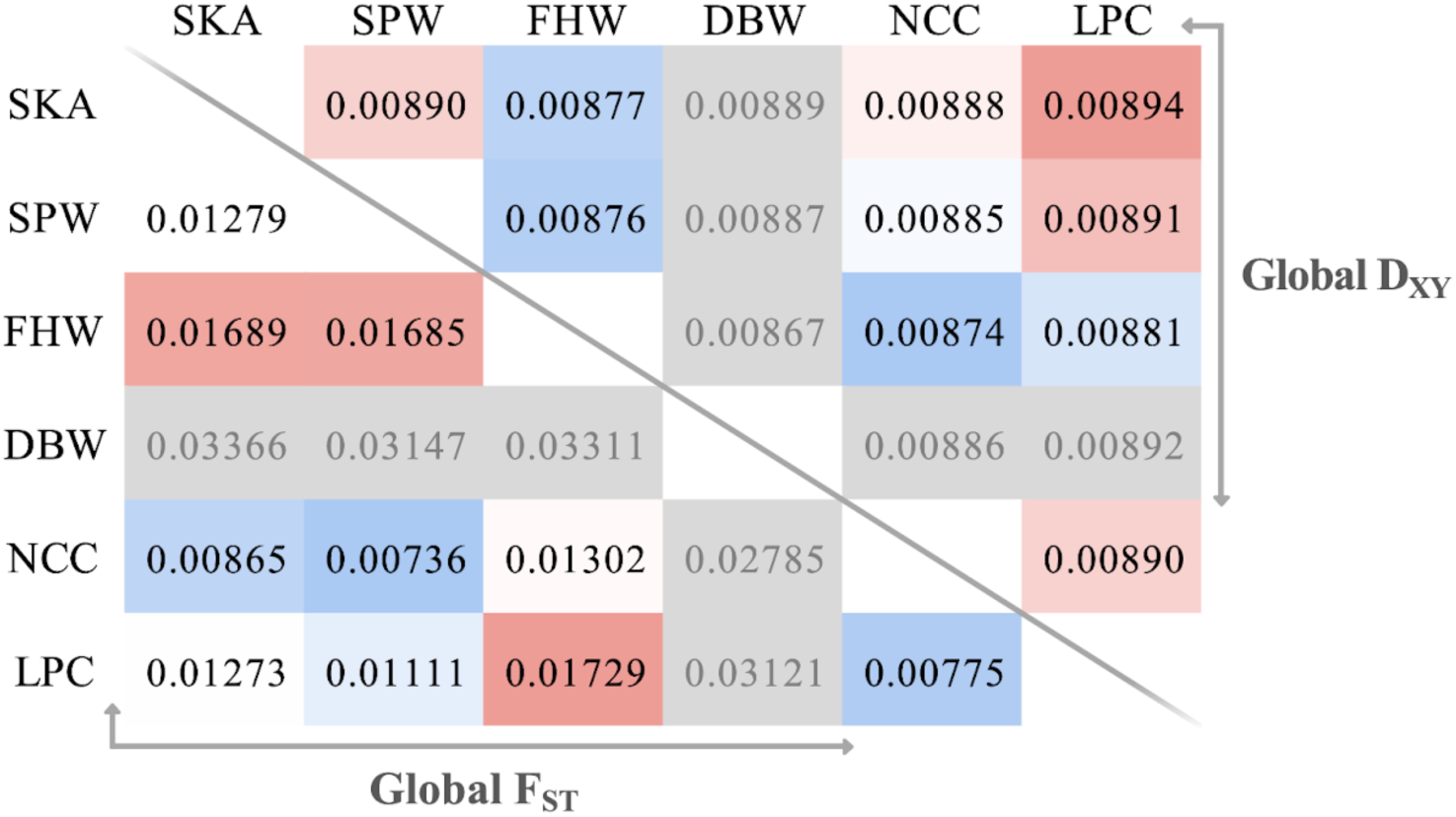
Heatmap displaying genome-wide averages for F_ST_ and D_XY_ by sampling location.

### Methods

Global π, F_ST_, and D_XY_ values were calculated on the quality-filtered all-sites vcf (Figure S2.2) for each sampling location (for π) or pair of sampling locations (F_ST_ and D_XY_) using pixy (Korunes & Samuk, 2021) in 15kb windows, then averaged across all windows along the genome. Specifically, estimates of π reflected the average per site nucleotide diversity, weighted by the number of genotyped samples at that site in that population for each 15kb window (Korunes & Samuk, 2021). Nucleotide divergence between populations (D_XY_) was calculated as the average nucleotide divergence at each site within each genomic window (Korunes & Samuk, 2021). Finally, estimates of global F_ST_ were generated for each pair of sampling locations (treated as populations) using the average Weir and Cockerham’s F_ST_ per SNP (as opposed to site) for each 15kb window (Korunes & Samuk, 2021). Notably, 15kb windows with negative F_ST_ values were converted to an F_ST_ of zero prior to calculating the average across all windows.

## Supplement S5. Population structure

**Figure S5.1.**
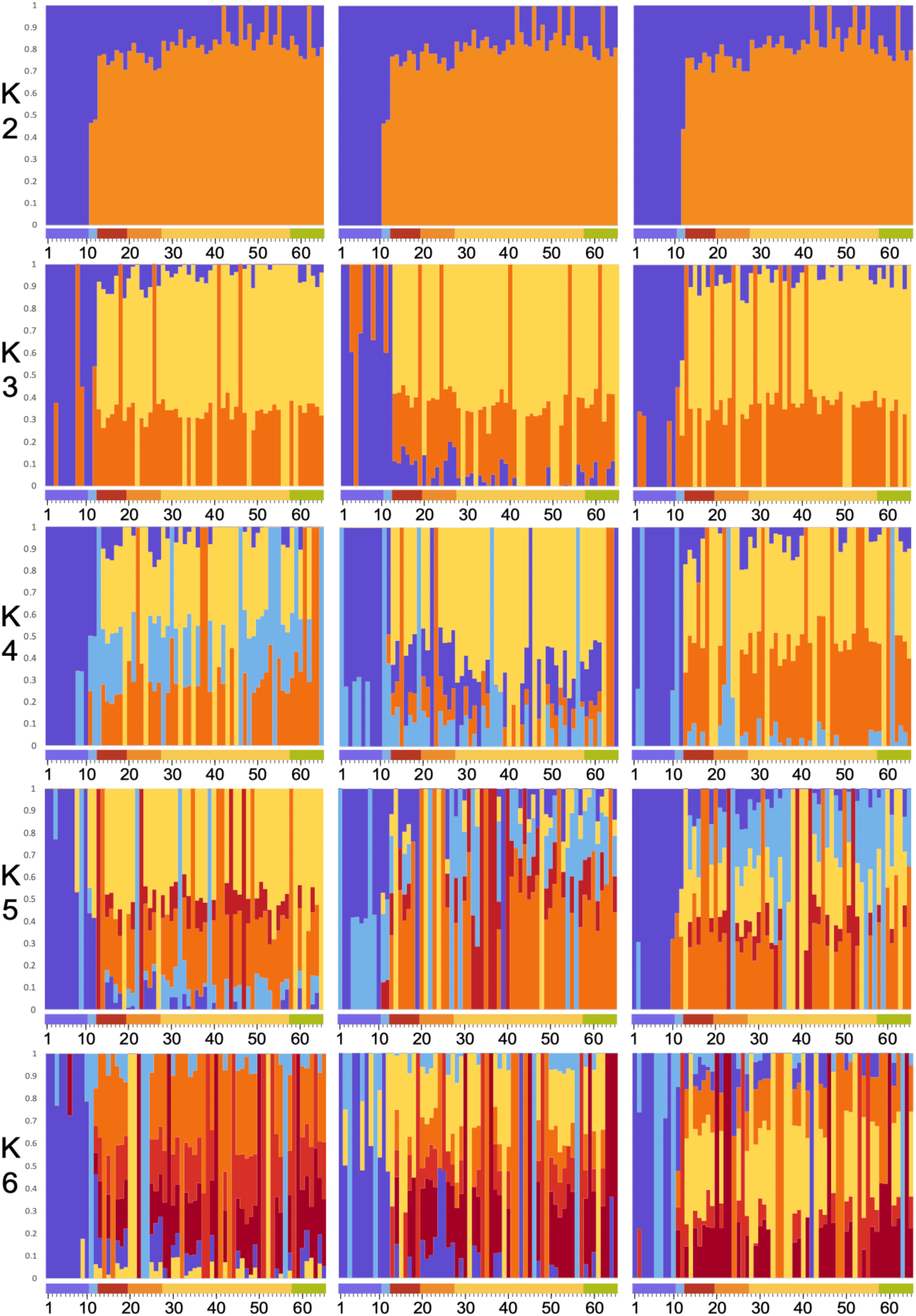
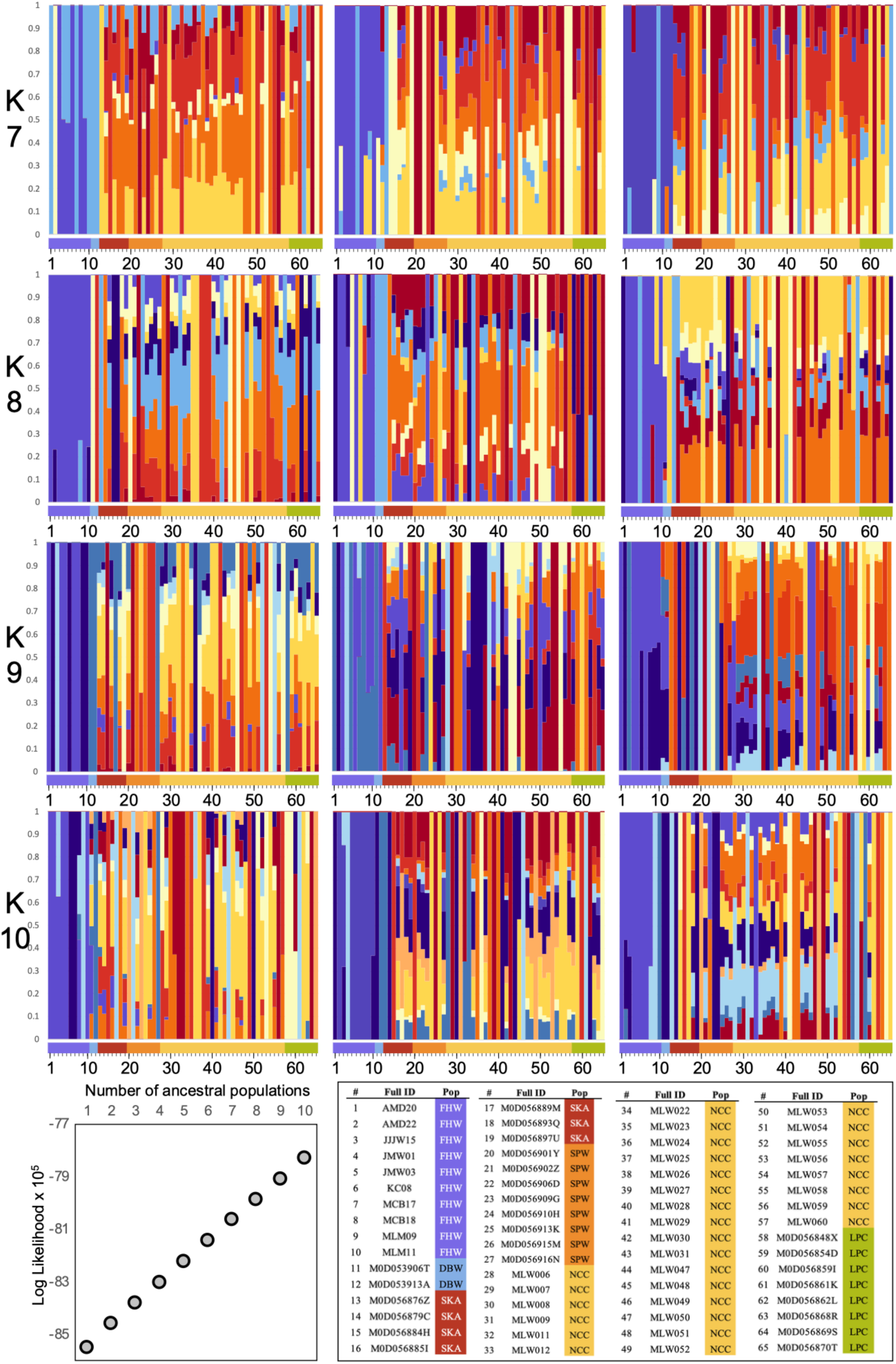
*(previous two pages)* **ADMIXTURE results on the neutral SNP panel for K=2 through K=10 run in triplicate.** Samples are ordered by location along x-axes and temporarily labeled (#), defined in the bottom-left panel on page 2. Bottom-right panel on page 2 gives log likelihood estimates for each value of K (number of ancestral populations) averaged across three replicates.

**Figure S5.2.**
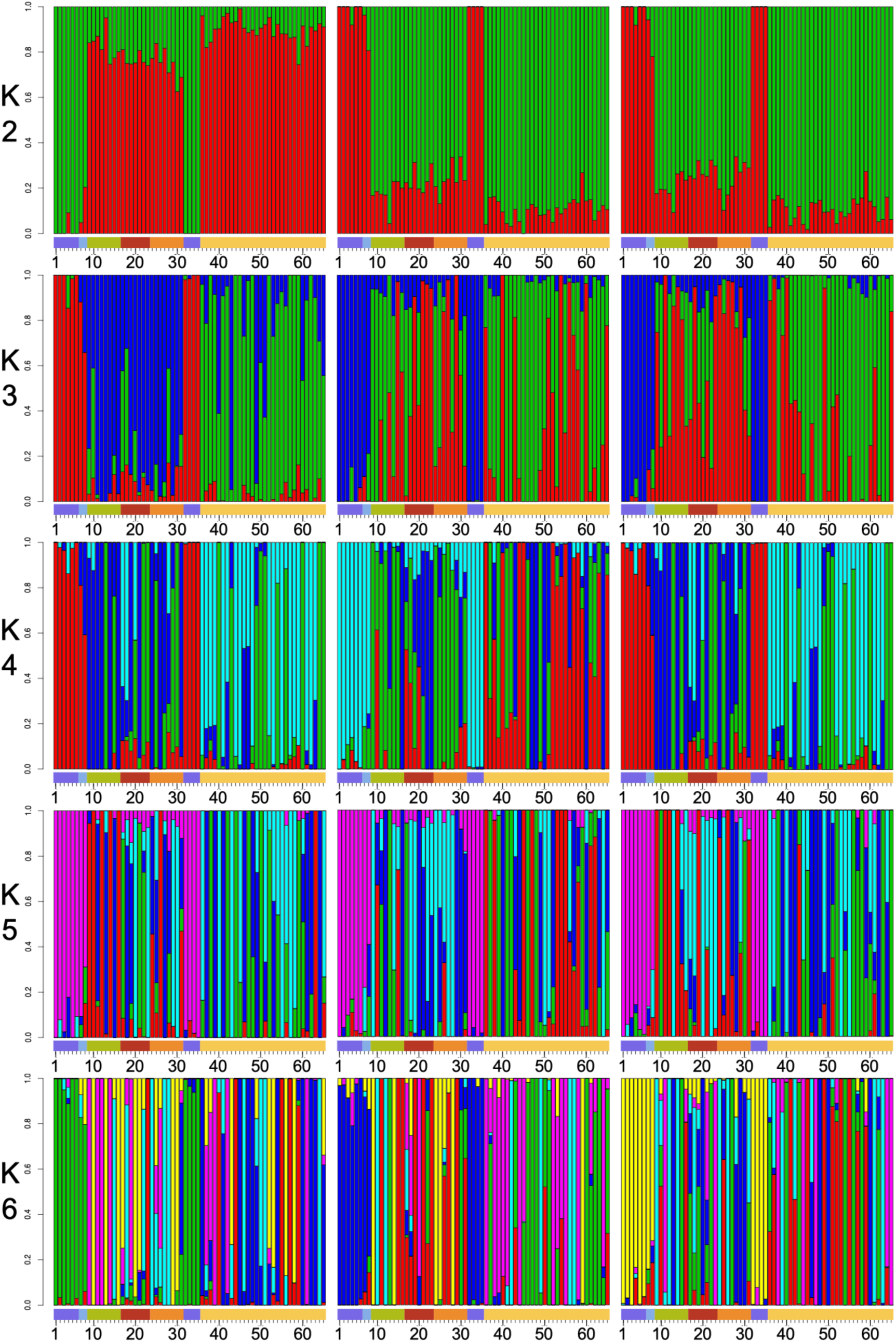
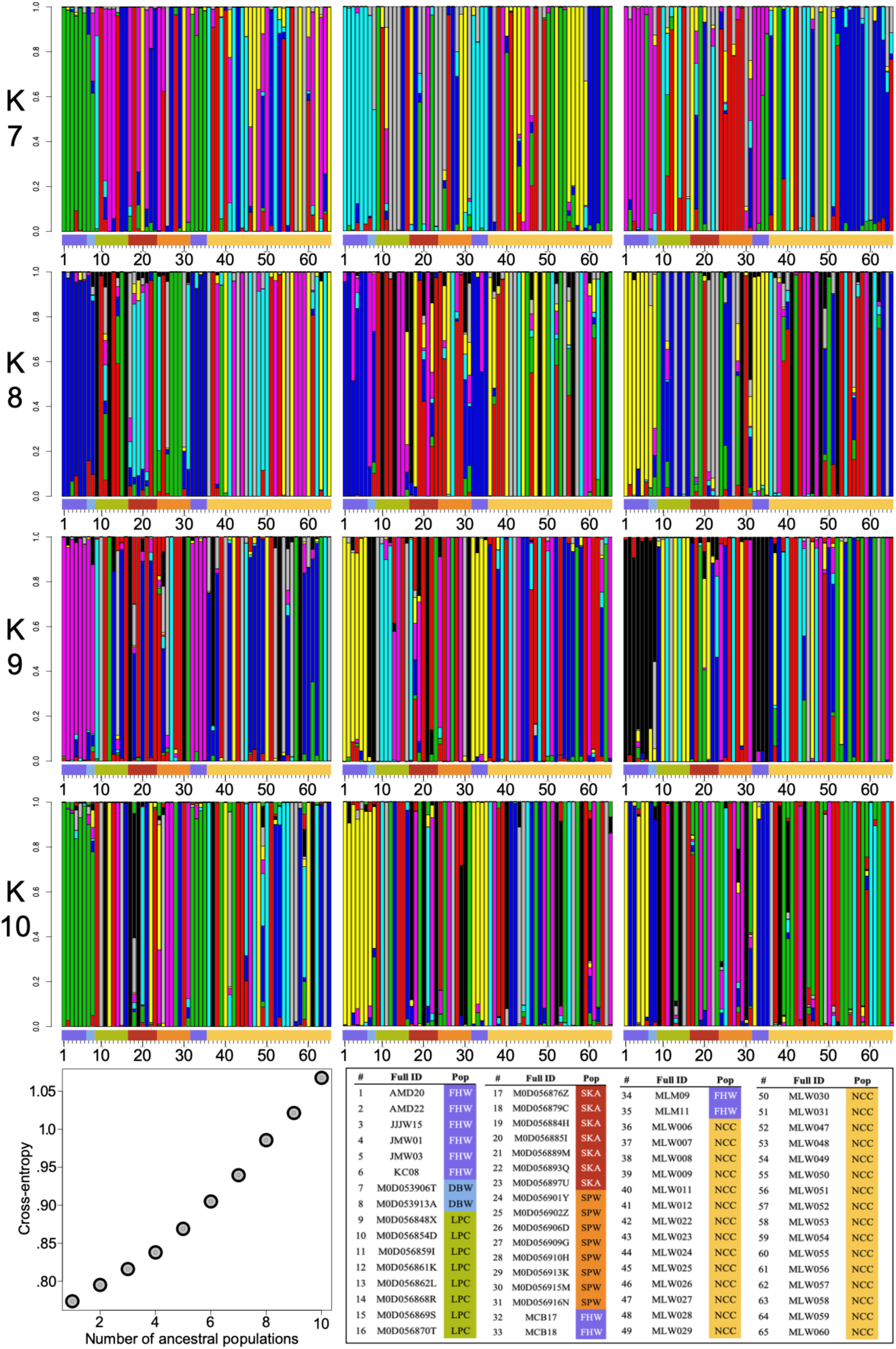
*(previous two pages)* **sNMF results on the neutral SNP panel for K=2 through K=10 run in triplicate.** Samples are ordered alphabetically along x-axes (with color bars added to indicate sampling location) and temporarily labeled (#), then later defined in the bottom-left panel on page 2. Bottom-right panel on page 2 gives sNMF (sparse Nonnegative Matrix Factorization) cross entropy estimates for each value of K (number of ancestral populations).

**Figure S5.3.**
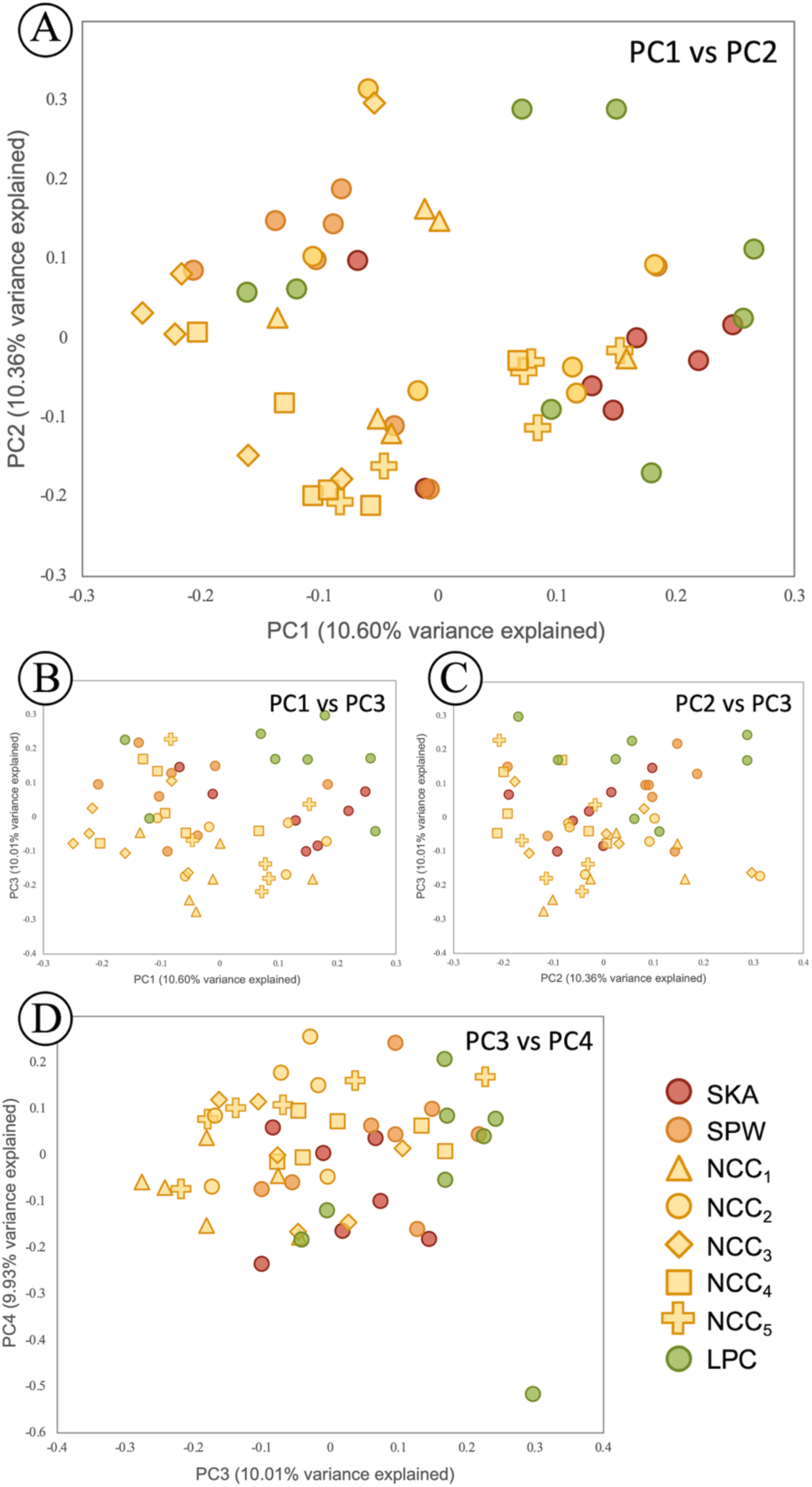
Principal components analysis (PCA) scatterplot for outer coast sites only, uniquely filtered (see main text methods), with North Central California (NCC) sites parsed into subsampled regions (see main text Figure 1). (**A**) PC1 plotted against PC2. (**B**) PC1 plotted against PC3. (**C**) PC2 plotted against PC3. (**D**) PC3 plotted against PC4.

### Extended ADMIXTURE results for K=2 through K=10 in triplicate (on neutral SNP panel)

### Extended sNMF results for K=2 through K=10 in triplicate (on neutral SNP panel)

### Outer coast only PCAs with subset-specific SNP filtering (but not neutral sites only)

## Supplement S6. Isolation by distance

**Figure S6.1.**
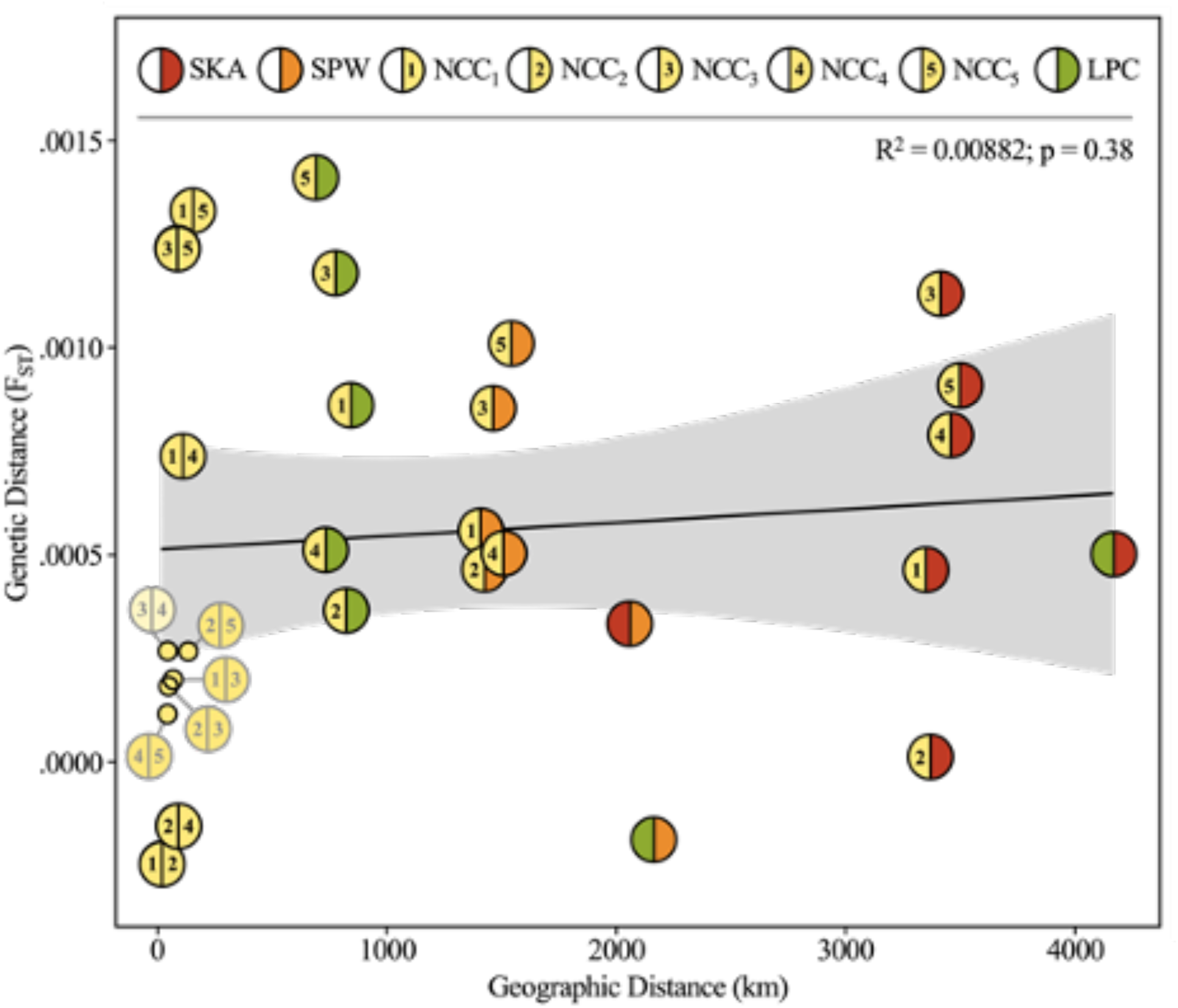
Isolation-by-distance analysis among stars collected from outer coast sites. Mantel test assessing isolation-by-distance (IBD) among sites in the OC population by correlating geographic distance (in km) to genetic distance (F_ST_), calculated using the neutral SNP panel, filtered to include only outer coast sites (SKA, SPW, NCC, and LPC). Mantel statistic is based on Pearson’s product-moment correlation and 999 permutations. Equation of the line: y = 0.00051 + 3.3e^-11^x.

## Supplement S7. Mitochondrial diversity

### Mitochondrial diversity among our samples

The location of the mitochondrion in the unpublished version 2 genome was determined by querying a subset of the shortest genome scaffolds approximating the reported length of the *P. ochraceus* mitochondrion at 16,376 bases (+/- 25%). These results were then BLASTed (Altschul et al., 1990) against the published (i.e., version 1) mitochondrion (genome accession number MH713001.1). The scaffold that best matched this sequence (“Scaffold_122 1_contigs length_16217”), at 99% similarity and a slightly larger size of 16,609 bp, was concluded to be the mitochondrion.

For direct comparison with the genetic analyses performed in Harley et al. (2006), we qualitatively assessed variability within the cytochrome oxidase c subunit I (COI) gene region by using CLUSTAL W (Thompson et al., 1994) to first map the 1554 bp of the published COI sequence for *P. ochraceus* (Gene ID: 40487529) to the mtDNA consensus sequence derived from mapping our reads to the version 2 *P. ochraceus* genome (Hu et al., *in prep*) in order to identify the region of interest.

**Figure S7.1.**
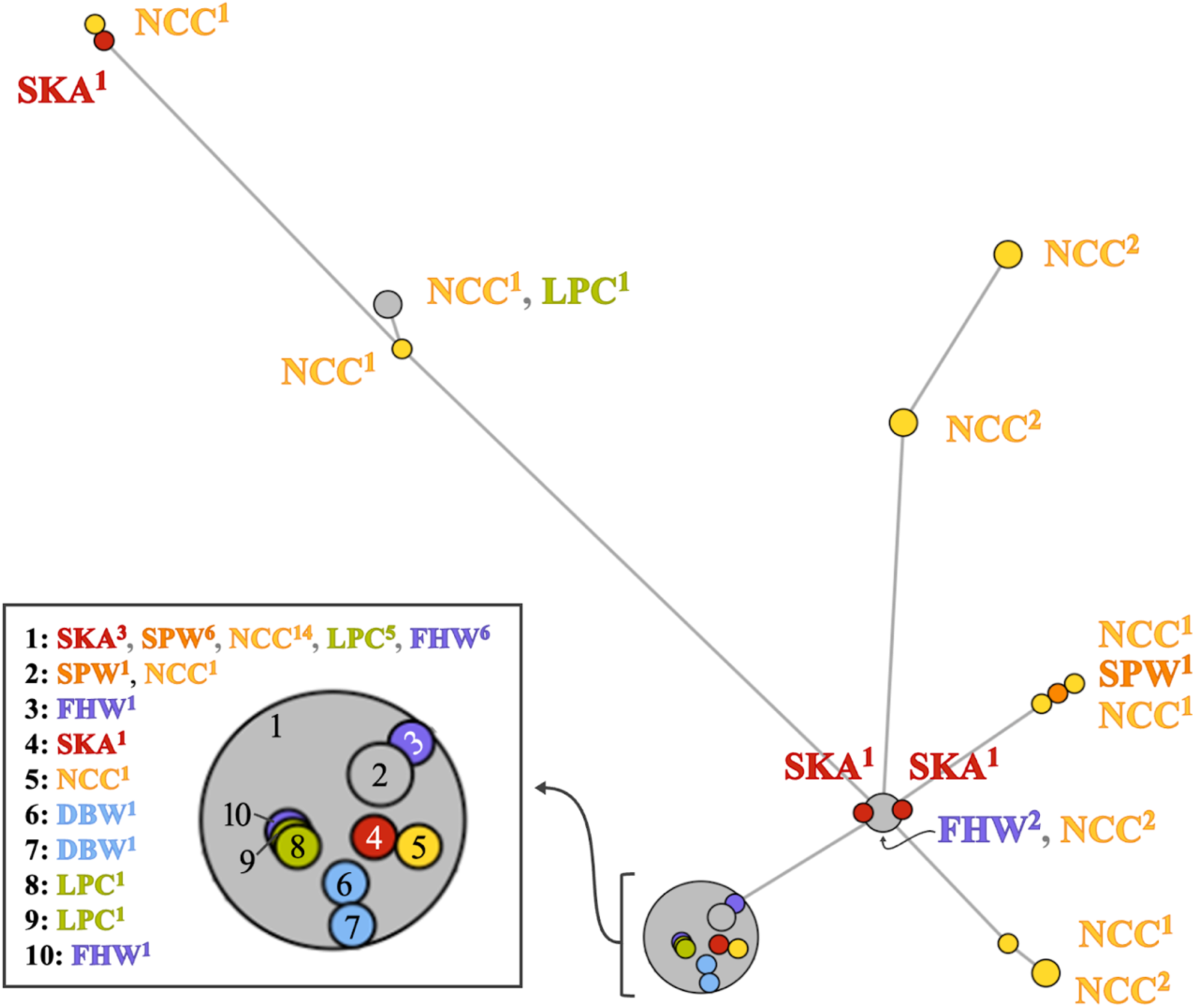
Haplotype network analysis for *P. ochraceus* mitochondrial genome. Size of the node bubble represents the total number of individuals comprising a haplotype, while site abbreviation superscripts represents the number of individuals collected from a given sampling location mapping to the labelled node.

### Comparative analysis with previous reports

In comparing our results with previous studies, two results appear to be consistent. (1) In Harley et al. (2006), a 543-bp region of COI presented a single very common haplotype (circled “A” in “Figure 8” from Harley et al., 2006; reprinted in Figure S7.2A) with a large number of singleton mutations related to it, a small group of sequences that were roughly 1.5% divergent, and some discussion of a ‘private haplotype’ (circled “G” in reprinted Figure S7.2A) that was only observed in Dabob Bay, Washington. (**2**) In Marko et al. (2010), the same 543-bp region of COI presented the same genealogical pattern overall (isolated haplotype network from Marko et al., 2010; reprinted in Figure S7.2B), with one mitotype at Cordova, Alaska being identical to the common haplotype at Dabob Bay and the other representing the less common divergent sequences shown in Figure S7.2A reprinted from Harley et al. (2006) Figure 8.

**Figure S7.2.**
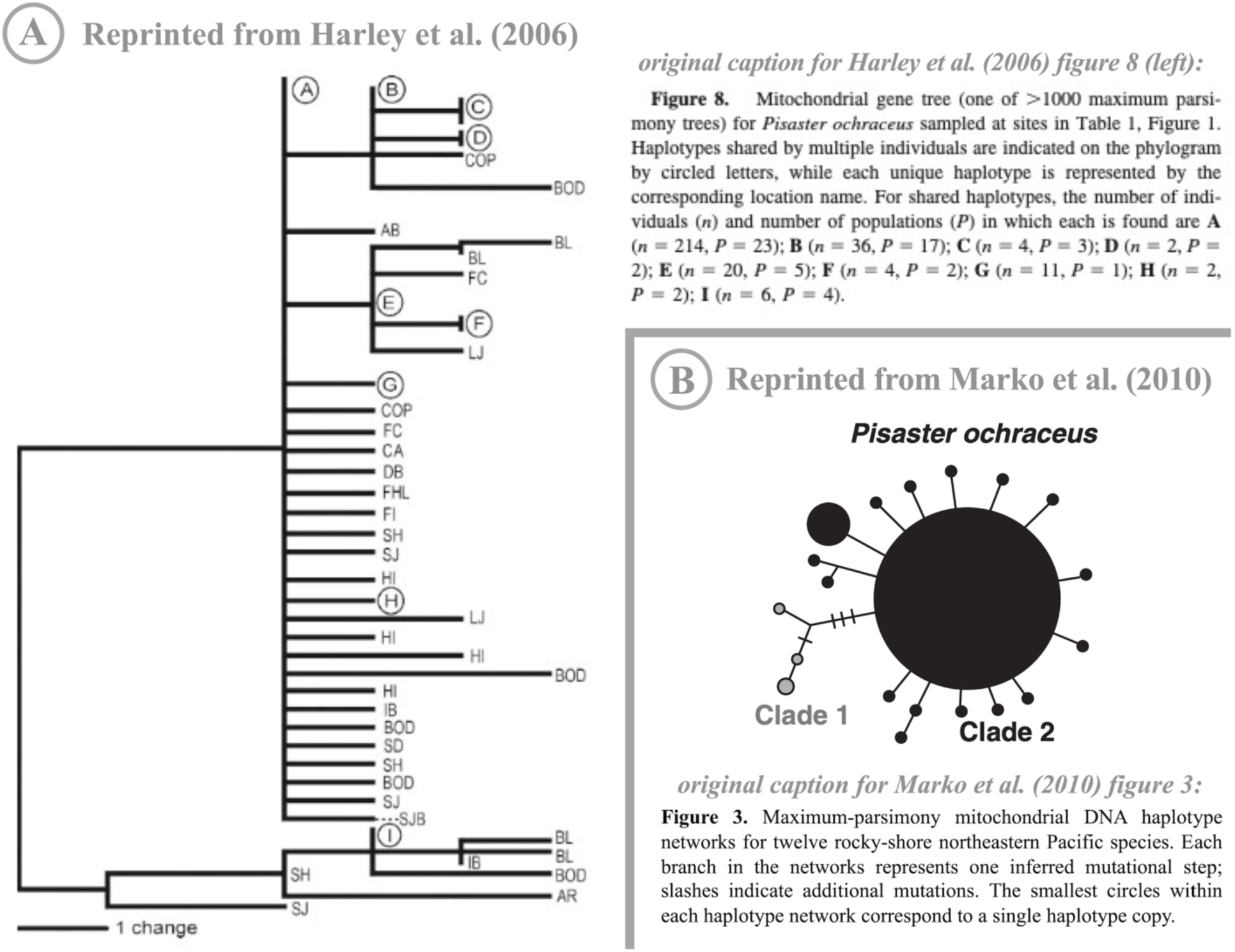
Reprinted figures displaying relationships among *P. ochraceus* mitochondrial haplotypes in (A) Harley et al. (2006) and (B) Marko et al. (2010) at a 543-bp region of the cytochrome c oxidase subunit I (COI) gene. Original figure labels and captions given within figure. (A) represents the entire figure 8 from Harley et al. (2006) while (B) represents a simplified version of figure 3 from Marko et al. (2010), cropped to include only the relevant species’ (*Pisaster ochraceus*) haplotype network.

When sequences from Dabob Bay (Harley et al., 2006; S7.2A) and Cordova (Marko et al., 2010; S7.2B) are aligned with the COI haplotypes generated from this study (Figure S7.3), the same pattern is recovered: a small frequency of haplotypes are relatively divergent from the most common haplotype (here amplified slightly because of an alignment issue JPW is having with the assembly breakpoint in a VCF - leads to fewer nucleotides in the overall alignment presented). In both Harley et al. (2006) and Marko et al., (2010) (shown in pink and blue, respectively; Figure S7.3), the private or distant mitotype present in those noted locations was not associated with an overall spatial pattern and the elevated distance was excluded from further consideration. We find overall consilience among the 3 studies and suggest that the sporadic appearance of one of the divergent or private haplotypes from one study at distinct locations in subsequent studies indicates more about the ‘chaotic genetic patchiness’ of demographic diversity in *Pisaster ochraceus* than a spatially divergent signal in the mitochondrial COI data (Figure S7.3).

**Figure S7.3.**
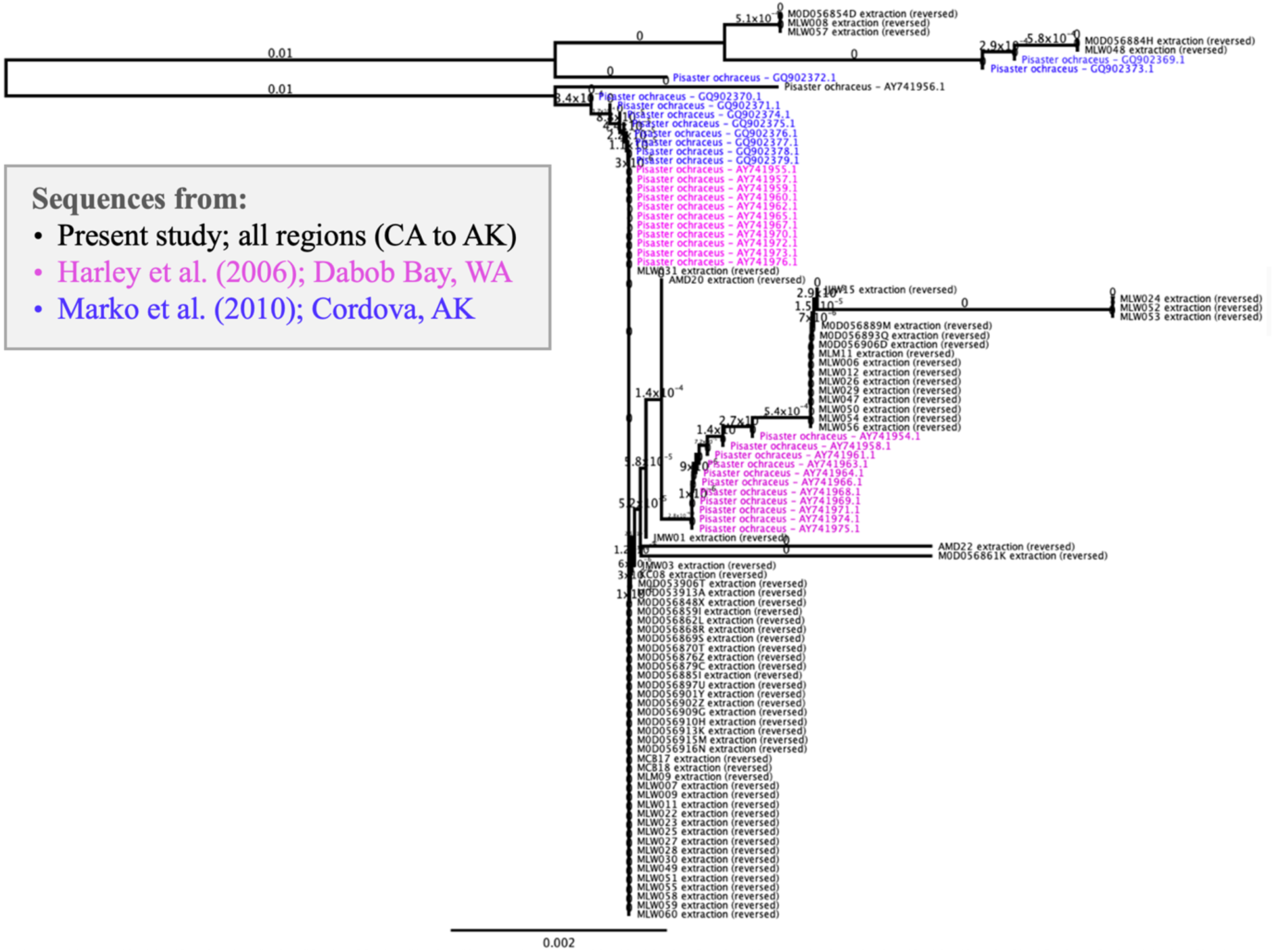
**Relationship between mitochondrial COI haplotypes generated from this study against those published in previous reports (Harley et al., 2006; Marko et al., 2010)**. Displayed as a neighbor-joining phylogram of aligned data from this study (marked in black) with Cordova (AK) sequences from Marko et al. (2010), marked in blue, and Dabob (WA) sequences from Harley et al. (2006), marked in pink.

## Supplement S8. Meadow plots: pairwise sampling locations

**Figure S8.1.**
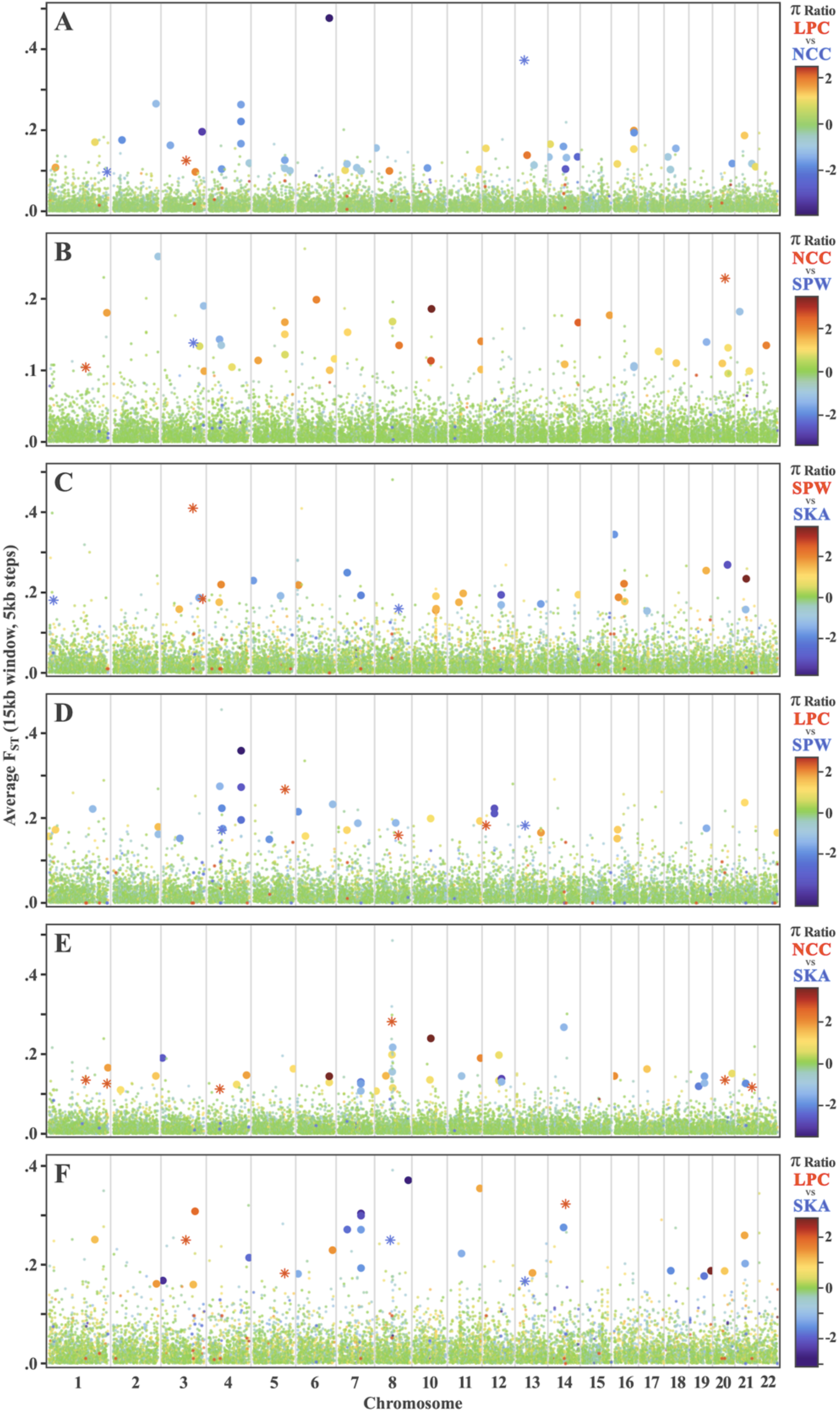
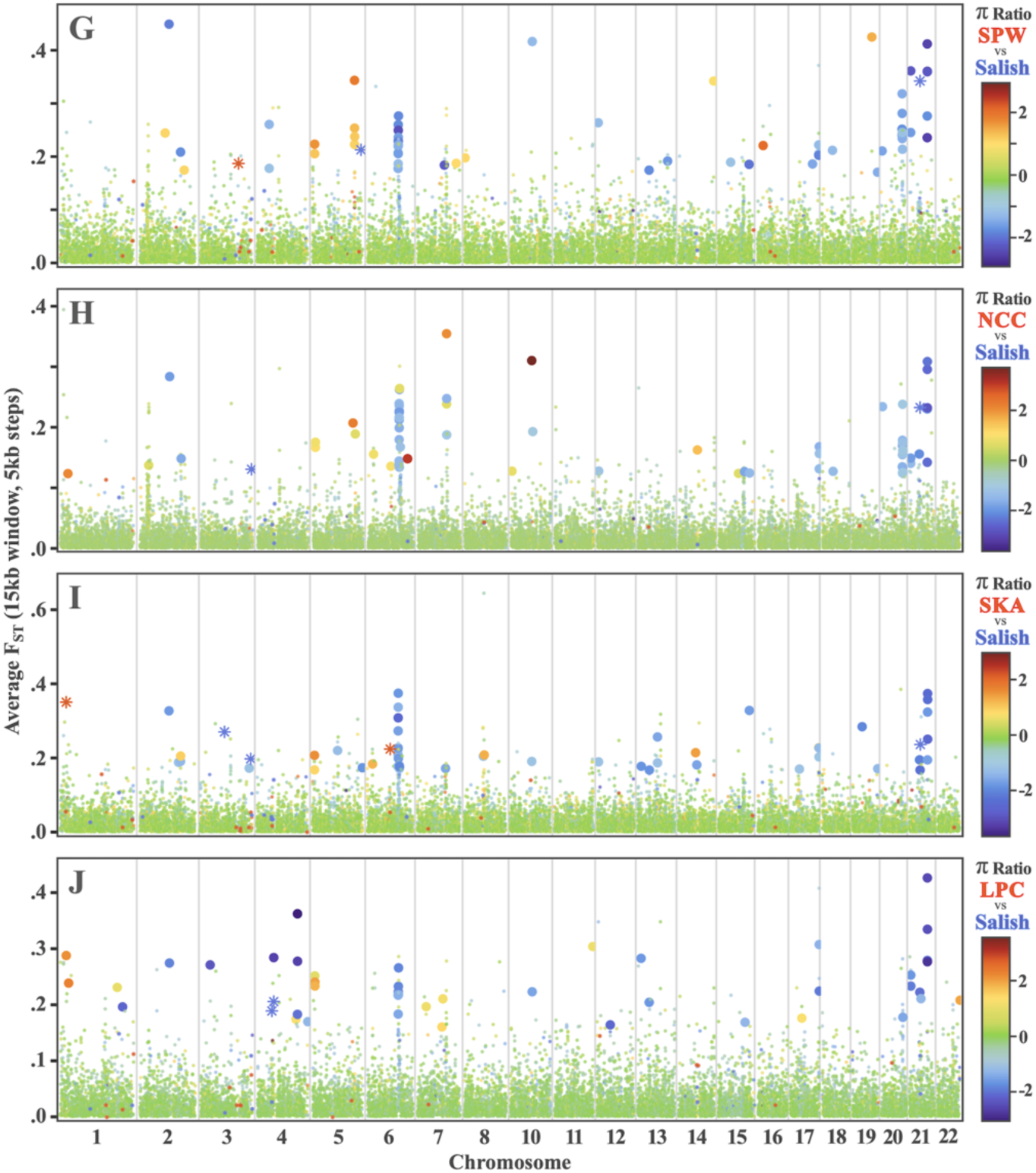
(starting on previous page) Full set of sampling location-based meadow plot results displayed as genomic regions of outlier F_ST_ and log_2_ π ratios across 15kb windows (with 5kb steps). The first panel (parts A-F) represents meadow plots wherein one OC site is being compared to another, while the second panel (parts G-J) features meadow plots that compare an OC site to the Salish Sea (i.e., FHW and DBW combined). Across all meadow plots, windows falling within both the top 1% of FST and the top or bottom 1% of log_2_ π ratios are considered outliers (defined using the same criteria pattern as described in Figure 4 caption), with datapoints representing outlier windows scaled up in size for emphasis. Finally, asterisk-shaped datapoints represent comparisons where one of the two populations under comparison had a π value of zero, resulting in a mathematical error calculation (+/-inf) for the log_2_ π ratio.

## Supplement S9. RDA Analysis (exemplar chromosomes)

**Figure S9.1.**
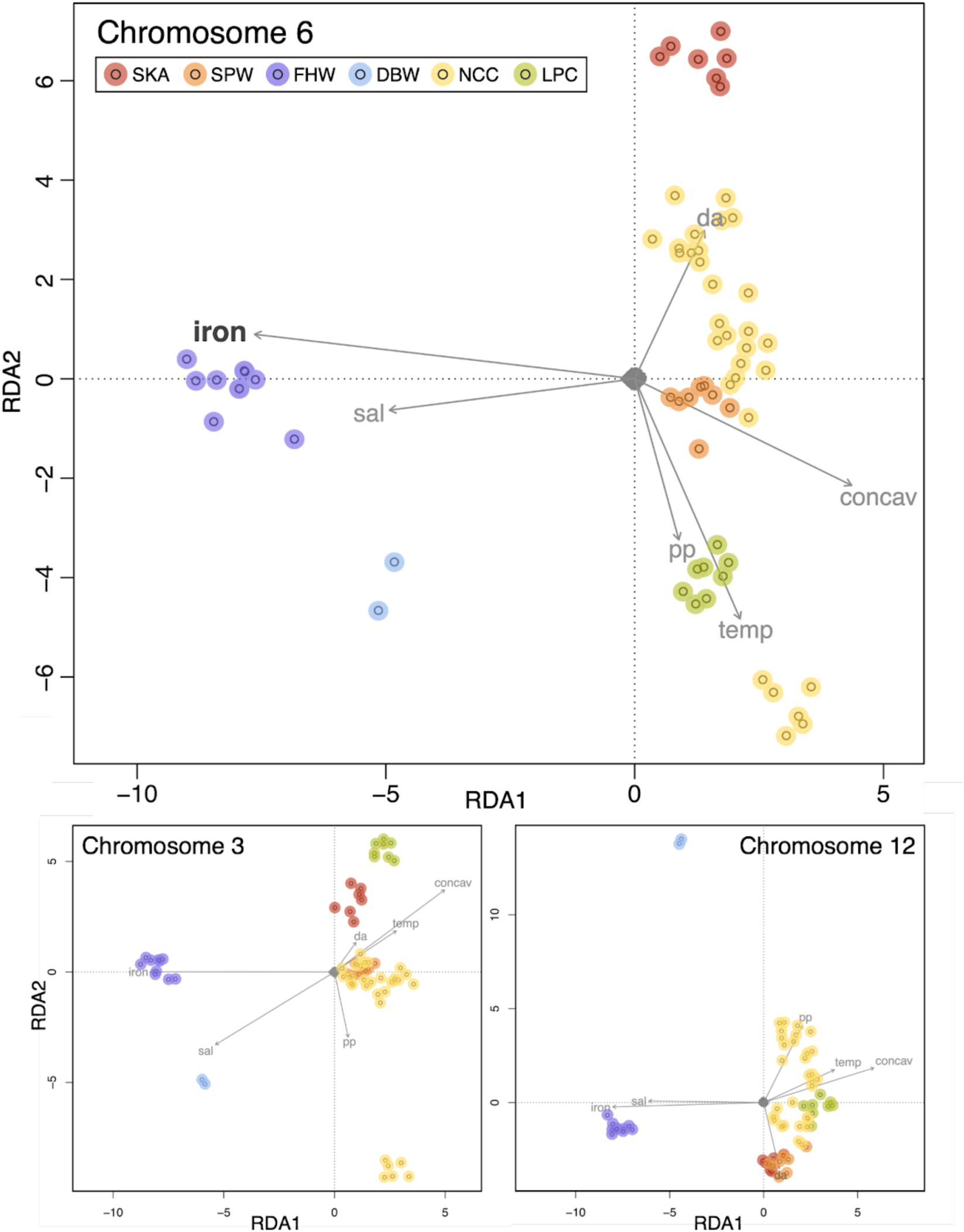
**Preliminary redundancy analysis (RDA) showing the relationships between genetic and environmental data**. Generated using full SNP panel (11,701,389 SNPs). Arrows indicate the direction and magnitude of environmental predictors.

### Methods

Environmental data obtained from sdmpredictors R package (Bosch & Fernandez, 2023; v0.2.15) following the developer’s protocol (http://lifewatch.github.io/sdmpredictors/). Input variables included: BO22_curvelmean_bdmean, BO22_dissoxmean_bdmean, BO22_ironmean_bdmean, BO22_lightbotmean_bdmean, BO22_ph, BO22_salinitymean_bdmean, BO22_tempmean_bdmean, BO22_temprange_bdmean, BO22_nitratemean_bdmean, BO22_ppmean_bdmean, BO22_curvelmax_bdmean, BO22_damean, BO22_calcite, and BO22_parmean.

## Supplement S10 supports Box 1. Community genomic transitions in the Strait of Juan de Fuca

This is where we summarize the data points in Box 1 for divergence information and larval information, thus putting the references for that figure here. All information currently is in <u>synthesis figure document</u>.

### Fishes

*Ophiodon elongatus*: Marko et al. (2007) only uses mitochondrial sequence data and there is a relatively large separation between the Salish samples and those along the outer coast.

Nevertheless, discusses distinction and lists ɸST as 0.025 between Salish and California samples in Tables 3 and 4 and Discussion. Describes PLD as “2.5 months” (70 days).

*Sebastes caurinus*: Buonaccorsi et al. (2002) use microsatellite data to identify distinct population in Salish Sea with F_ST_ 0.087 to Pacific coast samples. PLD claimed in paper as “2-3 months” so 70 days represented in Box 1.

*Sebastes auriculatus*: Buonaccorsi et al. (2005) use microsatellite data to identify distinct population in Salish Sea with F_ST_ 0.09 to Pacific coast samples. PLD claimed in paper as “2-3 months” so 70 days represented in Box 1; Kendall 1989 suggests PLD of 62 days.

*Gadus macrocephalus*: Drinan et al. (2018) compare *G. macrocephalus* from sites in the Salish sea to one at the opening of the strait of Juan de Fuca (F_ST_ 0.0148) and to an open ocean site offshore (F_ST_ 0.0187, value used in Box 1). Cunningham et al. (2009) identify a regional F_CT_ of 0.0094 between Salish and the outer coast (Table 4). References in Cunningham et al. (2009) lead to Hinckley et al. (2019) which suggests 70-75 days larval/juvenile duration.

*Sebastes ruberrimus*: Siegle et al. (2013) sample only a single “Strait of Georgia” site but find estimated F_ST_ 0.0172 as mean between that site and the nearest outer coast locations and argue for separation of this region. The larval duration is estimated at 60 days (COSEWIC 2008).

*Merluccius productus*: Longo et al. (2024) argue for a clear separation of the Salish populations and an F_ST_ between them and outer coast of 0.0104. Iwamoto et al. (2015) includes a contrast between Salish and California with mean F_ST_ of 0.035. PLD determined from Hollowed (1992).

### Mollusks

*Clinocardium nuttalli*: Dimond et al. (2022) identify clear divergence of Salish Sea populations of *Clinocardium* from open coast, with median F_ST_ 0.023 for this contrast. Pelagic larval duration (PLD) was listed as 10 days in this source.

*Ostrea lurida*: Silliman (2019) identifies clear divergence of *Ostrea* between Salish Sea and outer coast samples. Appendix C identifies F_ST_ of 0.108 and PLD is listed as “1-8 weeks” and is represented at 28 days in Box 1.

### Arthropods

*Cancer magister*: Jackson & O’Malley (2017) shows that *C. magister* exhibits very low F_ST_ (0.002) between the Salish and Outer Coast but this value is significant and much larger than other pairwise comparisons in their data. Jackson & O’Malley suggest PLD of “2-3 months” but Lough (1975) suggests 128 days.

*Balanus glandula*: Wares et al. (2021) provides RADseq data that show no evidence of distinct separation of the Salish population sampled relative to outer coast samples. *Balanus* is thus not represented on Box 1 beyond the PLD graph inset.

### Echinoderms

*Pisaster ochraceus*: see this study. PLD minimum (76 days) used in Box 1 as Strathmann (1978) argues “the shorter periods of larval development [observed in lab] are probably close to the usual pelagic period in nature.” We further modified this range using the data from Pia et al. (2012) to represent a more likely time to metamorphosis of 44-51 days.

*Apostichopus californicus*: Lowell et al. (2023) provides evidence that is more consistent with isolation by distance rather than a clear separation into distinct genotypic groups. *Apostichopus* is thus not represented on Box 1 beyond the PLD graph inset.

### Mammals

There is growing consensus that the social, behavioral, and genomic distinctions in the killer whale *Orcinus* (Moura et al., 2014) merit taxonomic recognition (Morin et al., 2024). However, as a species that does not have pelagic larval dispersal as part of its life history, *Orcinus* is not directly represented on Box 1. The current map used in this figure is from Shields et al. (2018) mapping the distribution of *Orcinus* in the Salish Sea.

### Algae

*Nereocystis luetkeana*: Gierke et al. (2023) show quite distinct genomic diversity in bull kelp in the Salish but use a general linear model and other statistics to argue that it is primarily an isolation-by-distance pattern. *Nereocystis* is thus not represented on Box 1.

## Notes

https://doi.org/10.26008/1912/bco-dmo.934772.1

### Competing Interest Statement

The authors have declared no competing interest.

## References

Alexander, D. H., Novembre, J., & Lange, K. (2009). Fast model-based estimation of ancestry in unrelated individuals. Genome Research, 19(9), 1655–1664.

Andersson, L., Ryman, N., Rosenberg, R., & Ståhl, G. (1981). Genetic variability in Atlantic herring (*Clupea harengus harengus*): description of protein loci and population data. Hereditas, 95(1), 69–7.

Andrews, C. A. (2010). Natural selection, genetic drift, and gene flow do not act in isolation in natural populations. Nature Education Knowledge, 3(10), 5.

Aquino, C. A., Besemer, R. M., DeRito, C. M., Kocian, J., Porter, I. R., Raimondi, P. T., … & Hewson, I. (2021). Evidence that microorganisms at the animal-water interface drive sea star wasting disease. Frontiers in Microbiology, 11, 610009.

Barrett-Lennard, L.G. and Ellis, G.M. (2001) *Population structure and genetic variability in northeastern Pacific Killer Whales: Towards an assessment of population viability*. Pacific Biological Station, Nanaimo, British Columbia: Canadian Science Advisory Secretariat.

Barry, J. P., Baxter, C. H., Sagarin, R. D., & Gilman, S. E. (1995). Climate-related, long-term faunal changes in a California rocky intertidal community. Science, 267(5198), 672–675.

Barth, J. A., Allen, S. E., Dever, E. P., Dewey, R. K., Evans, W., Feely, R. A., … & Wingard, C. E. (2019). Better regional ocean observing through cross-national cooperation: a case study from the Northeast Pacific. Frontiers in Marine Science, 6, 93.

Bashevkin, S. M., Lee, D., Driver, P., Carrington, E., & George, S. B. (2016). Prior exposure to low salinity affects the vertical distribution of *Pisaster ochraceus* (Echinodermata: Asteroidea) larvae in haloclines. Marine Ecology Progress Series, 542, 123–140.

Bashevkin, S.M., C.D. Dibble, R.P. Dunn, J.A. Hollarsmith, G. Ng, E.V. Satterthwaite, S.G. Morgan. 2020. Larval dispersal in a changing ocean with an emphasis on upwelling regions. Ecosphere 11: e03015.

Bates, A. E., Pecl, G. T., Frusher, S., Hobday, A. J., Wernberg, T., Smale, D. A., … & Watson, R. A. (2014). Defining and observing stages of climate-mediated range shifts in marine systems. Global Environmental Change, 26, 27–38.

Bay, R. A., Taylor, E. B., & Schluter, D. (2019). Parallel introgression and selection on introduced alleles in a native species. Molecular Ecology, 28(11), 2802–2813.

Benestan, L., Quinn, B., Maaroufi, H., Laporte, M., Clark, F., & Greenwood, S. et al. (2016). Seascape genomics provides evidence for thermal adaptation and current-mediated population structure in American lobster (*Homarus americanus*). Molecular Ecology, 25(20), 5073–5092. 10.1111/mec.13811

Berg, P., Jentoft, S., Star, B., Ring, K., Knutsen, H., & Lien, S. et al. (2015). Adaptation to Low Salinity Promotes Genomic Divergence in Atlantic Cod (*Gadus morhua L*.). Genome Biology and Evolution, 7(6), 1644–1663. 10.1093/gbe/evv093

Blanchette, C. A., Richards, D. V., Engle, J. M., Broitman, B. R., & Gaines, S. D. (2005). Regime shifts, community change and population booms of keystone predators at the Channel Islands. In Proceedings of the California Islands Symposium (Vol. 6, pp. 435–441).

Blanchette, C.A., Helmuth, B., & Gaines, S.D. (2007). Spatial patterns of growth in the mussel, *Mytilus californianus*, across a major oceanographic and biogeographic boundary at Point Conception, California, USA. Journal of Experimental Marine Biology and Ecology, 340(2), 126–148. 10.1016/j.jembe.2006.09.022

Bohonak, A. J. (1999). Dispersal, gene flow, and population structure. The Quarterly Review of Biology, 74(1), 21–45.

Booth, D. B., Troost, K. G., Clague, J. J., & Waitt, R. B. (2003). The Cordilleran ice sheet. Developments in Quaternary Sciences, 1, 17–43.

Briggs, J. C., & Bowen, B. W. (2013). Marine shelf habitat: biogeography and evolution. Journal of Biogeography, 40(6), 1023–1035.

Briggs, J. C., & Bowen, B. W. (2012). A realignment of marine biogeographic provinces with particular reference to fish distributions. Journal of Biogeography, 39(1), 12–30.

Buonaccorsi, V. P., Kimbrell, C. A., Lynn, E. A., & Vetter, R. D. (2002). Population structure of copper rockfish (*Sebastes caurinus*) reflects postglacial colonization and contemporary patterns of larval dispersal. Canadian Journal of Fisheries and Aquatic Sciences, 59(8), 1374–1384.

Buonaccorsi, V. P., Kimbrell, C. A., Lynn, E. A., & Vetter, R. D. (2005). Limited realized dispersal and introgressive hybridization influence genetic structure and conservation strategies for brown rockfish, Sebastes auriculatus. Conservation Genetics, 6, 697–713.

Byers, J. E., & Pringle, J. M. (2006). Going against the flow: retention, range limits and invasions in advective environments. Marine Ecology Progress Series, 313, 27–41.

Chai, Z., Xin, J., Zhang, C., Dawayangla, Luosang, Zhang, Q., Pingcuozhandui, Li, C., Zhu, Y., Cao, H., Wang, H., Han, J., Ji, Q., & Zhong, J. (2020). Whole-genome resequencing provides insights into the evolution and divergence of the native domestic yaks of the Qinghai–Tibet plateau. BMC Evolutionary Biology, 20(1). 10.1186/s12862-020-01702-8

Chandler, V. K., & Wares, J. P. (2017). RNA expression and disease tolerance are associated with a “keystone mutation” in the ochre sea star *Pisaster ochraceus*. PeerJ, 5, e3696

Chase, Z., Strutton, P. G., & Hales, B. (2007). Iron links river runoff and shelf width to phytoplankton biomass along the US West Coast. Geophysical Research Letters, 34(4).

Chen, Y., Wu, X., Wang, J., Hou, Y., Liu, Y., Wang, B., … & Yin, Z. (2022). Detection of selection signatures in Anqing six-end-white pigs based on resequencing data. Genes, 13(12), 2310.

Christensen, V., Coll, M., Piroddi, C., Steenbeek, J., Buszowski, J., & Pauly, D. (2014). A century of fish biomass decline in the ocean. Marine Ecology Progress Series, 512, 155–166.

Clague, J.J. (1989). Quaternary sea levels (Canadian Cordillera). In Quaternary geology of Canada and Greenland. Chap. 1. Geology of Canada. No. 1. Edited by R.J. Fulton. Geological Survey of Canada, Ottawa. pp. 43–47.

Clausen, R., & York, R. (2008). Economic growth and marine biodiversity: influence of human social structure on decline of marine trophic levels. Conservation Biology, 22(2), 458–466.

Cowen, R. K., & Sponaugle, S. (2009). Larval dispersal and marine population connectivity. Annual Review of Marine Science, 1, 443–466. 10.1146/annurev.marine.010908.163757

Cunningham, K. M., Canino, M. F., Spies, I. B., & Hauser, L. (2009). Genetic isolation by distance and localized fjord population structure in Pacific cod (*Gadus macrocephalus*): limited effective dispersal in the northeastern Pacific Ocean. Canadian Journal of Fisheries and Aquatic Sciences, 66(1), 153–166.

Danecek, P., Auton, A., Abecasis, G., Albers, C. A., Banks, E., DePristo, M. A., … & 1000 Genomes Project Analysis Group. (2011). The variant call format and VCFtools. Bioinformatics, 27(15), 2156–2158.

Dawson, M. N., Raskoff, K. A., & Jacobs, D. K. (1998). Field preservation of marine invertebrate tissue for DNA analyses. Molecular Marine Biology and Biotechnology, 7(2), 145–152.

Dawson, M.N. (2001). Phylogeography in coastal marine animals: a solution from California? Journal of Biogeography, 28(6), 723–736. 10.1046/j.1365-2699.2001.00572.x

Dawson, M. N. (2014). Natural experiments and meta-analyses in comparative phylogeography. Journal of Biogeography, 41(1), 52–65.

Dawson, M. N., Barber, P. H., González-Guzmán, L. I., Toonen, R. J., Dugan, J. E., & Grosberg, R. K. (2011). Phylogeography of *Emerita analoga* (Crustacea, Decapoda, Hippidae), an eastern Pacific Ocean sand crab with long-lived pelagic larvae. Journal of Biogeography, 38(8), 1600–1612.

Dawson, M. N., Hays, C. G., Grosberg, R. K., & Raimondi, P. T. (2014). Dispersal potential and population genetic structure in the marine intertidal of the eastern North Pacific. Ecological Monographs, 84(3), 435–456.

Dawson, M. N., Duffin, P. J., Giakoumis, M., Schiebelhut, L. M., Beas-Luna, R., Bosley, K. L., … & Wares, J. P. (2023). A decade of death and other dynamics: deepening perspectives on the diversity and distribution of sea stars and wasting. The Biological Bulletin, 244(3), 139–219.

Dayan, D. I. (2020). Clinal adaptation in the marine environment. Population Genomics: Marine Organisms, 221-247.

DeBiasse, M. B., Schiebelhut, L. M., Escalona, M., Beraut, E., Fairbairn, C., Marimuthu, M. P., … & Dawson, M. N. (2022). A chromosome-level reference genome for the giant pink sea star, *Pisaster brevispinus*, a species severely impacted by wasting. Journal of Heredity, 113(6), 689–698.

DePristo, M. A., Banks, E., Poplin, R., Garimella, K. V., Maguire, J. R., Hartl, C., … & Daly, M. J. (2011). A framework for variation discovery and genotyping using next-generation DNA sequencing data. Nature Genetics, 43(5), 491–498.

Dietrich, M. R., Ankeny, R. A., Crowe, N., Green, S., & Leonelli, S. (2020). How to choose your research organism. Studies in History and Philosophy of Science Part C: Studies in History and Philosophy of Biological and Biomedical Sciences, 80, 101227.

Duffin, P., Martin, D. L., Furman, B. T., & Ross, C. (2021). Spatial patterns of *Thalassia testudinum* immune status and *Labyrinthula* spp. load implicate environmental quality and history as modulators of defense strategies and wasting disease in Florida Bay, United States. Frontiers in Plant Science, 12, 612947.

Duffin, P.J. (2024). Beast under burden: Characterizing range-wide genomic diversity and gene expression patterns in the keystone predator *Pisaster ochraceus* amid modern threats. Ph.D. Dissertation, University of Georgia

Dungan, M. L., Miller, T. E., & Thomson, D. A. (1982). Catastrophic decline of a top carnivore in the Gulf of California rocky intertidal zone. Science, 216(4549), 989–991.

Ebbesmeyer, C. C., & Barnes, C. A. (1980). Control of a fjord basin’s dynamics by tidal mixing in embracing sill zones. Estuarine and Coastal Marine Science, 11(3), 311–330.

Ebbesmeyer, C. C., Cannon, G. A., & Barnes, C. A. (1984). Synthesis of current measurements in Puget Sound, Washington. Volume 3: Circulation in Puget Sound: An interpretation based on historical records of currents (NOAA Technical Memorandum NOS OMS 5, pp. 1–73). U.S. Department of Commerce, National Oceanic and Atmospheric Administration.

Eisenlord, M. E., Groner, M. L., Yoshioka, R. M., Elliott, J., Maynard, J., Fradkin, S., … & Harvell, C. D. (2016). Ochre star mortality during the 2014 wasting disease epizootic: role of population size structure and temperature. Philosophical Transactions of the Royal Society B: Biological Sciences, 371(1689), 20150212.

Eldon, B., Riquet, F., Yearsley, J., Jollivet, D., & Broquet, T. (2016). Current hypotheses to explain genetic chaos under the sea. Current Zoology, 62(6), 551–566.

Ewers-Saucedo, C., J.M. Pringle, H.H. Sepúlveda, J.E. Byers, S.A. Navarrete, and J.P. Wares. 2016. The oceanic concordance of phylogeography and biogeography: a case study in *Notochthamalus*. Ecol. Evol. 6: 4403–4420.

Felsenstein, J. (1985). Phylogenies and the comparative method. The American Naturalist, 125(1), 1–15.

Feng, P., Zeng, T., Yang, H., Chen, G., Du, J., Chen, L., … & Lu, L. (2021). Whole-genome resequencing provides insights into the population structure and domestication signatures of ducks in eastern China. BMC Genomics, 22(1), 401.

Flower, Aquila, “Atlas of the Salish Sea” (2020). Salish Sea Maps. 2. https://cedar.wwu.edu/salish_maps/2

Forester, B. R., Lasky, J. R., Wagner, H. H., & Urban, D. L. (2018). Comparing methods for detecting multilocus adaptation with multivariate genotype–environment associations. Molecular Ecology, 27(9), 2215–2233.

Frichot, E., & François, O. (2015). LEA: An R package for landscape and ecological association studies. Methods in Ecology and Evolution, 6(8), 925–929.

Frichot, E., Mathieu, F., Trouillon, T., Bouchard, G., & François, O. (2014). Fast and efficient estimation of individual ancestry coefficients. Genetics, 196(4), 973–983.

Frontana-Uribe, S., J. de la Rosa-Vélez, L. Enríquez-Paredes, L.B. Ladah, and L. Sanvincente-Añorve. 2008. Lack of genetic evidence for the subspeciation of *Pisaster ochraceus* (Echinodermata: Asteroidea) in the northeastern Pacific Ocean. J. Mar. Biol. Assoc. UK. 88: 395–400.

Gierke, L., Coelho, N. C., Khangaonkar, T., Mumford, T., & Alberto, F. (2023). Range wide genetic differentiation in the bull kelp *Nereocystis luetkeana* with a seascape genetic focus on the Salish Sea. Frontiers in Marine Science, 10, 1275905.

González-Delgado, S., Lozano-Bilbao, E., Hardisson, A., Paz, S., Gonález-Weller, D., Rubio, C., & Gutiérrez, Á. J. (2024). Metal concentrations in echinoderms: Assessing bioindicator potential and ecological implications. Marine Pollution Bulletin, 205, 116619.

Gooding, R. A., Harley, C. D., & Tang, E. (2009). Elevated water temperature and carbon dioxide concentration increase the growth of a keystone echinoderm. Proceedings of the National Academy of Sciences, 106(23), 9316–9321.

Gruber, B., Unmack, P. J., Berry, O. F., & Georges, A. (2018). dartR: An R package to facilitate analysis of SNP data generated from reduced representation genome sequencing. Molecular Ecology Resources, 18(3), 691–699.

Guo, B., & Wares, J. P. (2017). Large-scale gene flow in the barnacle Jehlius cirratus and contrasts with other broadly-distributed taxa along the Chilean coast. PeerJ, 5, e2971.

Guo, T., Zhao, H., Yue, Y., Yang, Y., Wang, X., & Chen, Y. (2021). Selective sweeps uncovering the genetic basis of horn and adaptability traits on fine-wool sheep in China. Frontiers in Genetics, 12, 604235.

Harley, C., Pankey, M., Wares, J., Grosberg, R., & Wonham, M. (2006). Color polymorphism and genetic structure in the sea star *Pisaster ochraceus*. The Biological Bulletin, 211(3), 248–262. 10.2307/4134547

Held, M. B., & Harley, C. D. (2009). Responses to low salinity by the sea star *Pisaster ochraceus* from high-and low-salinity populations. Invertebrate Biology, 128(4), 381–390.

Herlinveaux, R. H., & Tully, J. P. (1961). Some oceanographic features of Juan de Fuca Strait. Journal of the Fisheries Board of Canada, 18(6), 1027–1071.

Hess, J. E., Campbell, N. R., Close, D. A., Docker, M. F., & Narum, S. R. (2013). Population genomics of Pacific lamprey: adaptive variation in a highly dispersive species. Molecular Ecology, 22(11), 2898–2916.

Hice, L. A., Duffy, T. A., Munch, S. B., & Conover, D. O. (2012). Spatial scale and divergent patterns of variation in adapted traits in the ocean. Ecology Letters, 15(6), 568–575.

Hodin, J., Heyland, A., Mercier, A., Pernet, B., Cohen, D. L., Hamel, J. F., … & George, S. B. (2019). Culturing echinoderm larvae through metamorphosis. Methods in Cell Biology, 150, 125–169.

Hoey, J. A., & Pinsky, M. L. (2018). Genomic signatures of environmental selection despite near-panmixia in summer flounder. Evolutionary Applications, 11(9), 1732–1747.

Jackson, T. M., & O’Malley, K. G. (2017). Comparing genetic connectivity among Dungeness crab (*Cancer magister*) inhabiting Puget Sound and coastal Washington. Marine Biology, 164, 1–12.

Jeffery, N., Stanley, R., Wringe, B., Guijarro-Sabaniel, J., Bourret, V., & Bernatchez, L. et al. (2017). Range-wide parallel climate-associated genomic clines in Atlantic salmon. Royal Society Open Science, 4(11), 171394. 10.1098/rsos.171394

Ji, L., Jordan, W. T., Shi, X., Hu, L., He, C., & Schmitz, R. J. (2018). TET-mediated epimutagenesis of the *Arabidopsis thaliana* methylome. Nature Communications, 9(1), 895.

Johnson, M. S., & Black, R. (1982). Chaotic genetic patchiness in an intertidal limpet, *Siphonaria* sp. Marine Biology, 70, 157–164. 10.1007/BF00397680

Jombart, T. (2008). adegenet: a R package for the multivariate analysis of genetic markers. Bioinformatics, 24(11), 1403–1405.

Jombart, T., Devillard, S., & Balloux, F. (2010). Discriminant analysis of principal components: a new method for the analysis of genetically structured populations. BMC Genetics, 11, 1–15.

Kardos, M., Zhang, Y., Parsons, K. M., Kang, H., Xu, X., Liu, X., … & Li, S. (2023). Inbreeding depression explains killer whale population dynamics. Nature Ecology & Evolution, 7(5), 675–686.

Kelly, M. W., Sanford, E., & Grosberg, R. K. (2012). Limited potential for adaptation to climate change in a broadly distributed marine crustacean. Proceedings of the Royal Society B: Biological Sciences, 279(1727), 349–356.

Kelly, R. P., & Palumbi, S. R. (2010). Genetic structure among 50 species of the northeastern Pacific rocky intertidal community. PLOS ONE, 5(1), e8594.

Khangaonkar, T., Nugraha, A., Xu, W., & Balaguru, K. (2019). Salish Sea response to global climate change, sea level rise, and future nutrient loads. Journal of Geophysical Research: Oceans, 124(6), 3876–3904.

Kimura, M. (1979). The neutral theory of molecular evolution. Scientific American, 241(5), 98–129.

Knaus, B. J., & Grünwald, N. J. (2017). vcfR: a package to manipulate and visualize variant call format data in R. Molecular Ecology Resources, 17(1), 44–53.

Kohl, W., McClure, T., & Miner, B. (2016). Decreased temperature facilitates short-term sea star wasting disease survival in the keystone intertidal sea star *Pisaster ochraceus*. PLOS ONE, 11(4), p. 0153670.

Korunes, K. L., & Samuk, K. (2021). pixy: Unbiased estimation of nucleotide diversity and divergence in the presence of missing data. Molecular Ecology Resources, 21(4), 1359–1368.

Krueger, F. (2015). Trim Galore!: A wrapper around Cutadapt and FastQC to consistently apply adapter and quality trimming to FastQ files, with extra functionality for RRBS data. *Babraham Institute*.

Lafferty, K. D., Harvell, C. D., Conrad, J. M., Friedman, C. S., Kent, M. L., Kuris, A. M., … & Saksida, S. M. (2015). Infectious diseases affect marine fisheries and aquaculture economics. Annual Review of Marine Science, 7(1), 471–496.

Lawrence, J. M. (Ed.). (2013). *Starfish: Biology and ecology of the Asteroidea*. JHU Press. Lawlor, J. A., & Arellano, S. M. (2020). Temperature and salinity, not acidification, predict near-future larval growth and larval habitat suitability of Olympia oysters in the Salish Sea. Scientific Reports, 10(1), 13787.

Li, H., & Durbin, R. (2009). Fast and accurate short read alignment with Burrows–Wheeler transform. Bioinformatics, 25(14), 1754–1760.

Li, H., Handsaker, B., Wysoker, A., Fennell, T., Ruan, J., Homer, N., … & 1000 Genome Project Data Processing Subgroup. (2009). The sequence alignment/map format and SAMtools. Bioinformatics, 25(16), 2078–2079.

Li, X., Yang, J. I., Shen, M., Xie, X. L., Liu, G. J., Xu, Y. X., … & Li, M. H. (2020). Whole-genome resequencing of wild and domestic sheep identifies genes associated with morphological and agronomic traits. Nature Communications, 11(1), 2815.

Liggins, L., Treml, E. A., & Riginos, C. (2020). Seascape genomics: contextualizing adaptive and neutral genomic variation in the ocean environment. Population Genomics: Marine Organisms, 171–218.

Lotterhos, K. E. (2019). The effect of neutral recombination variation on genome scans for selection. G3: Genes, Genomes, Genetics, 9(6), 1851–1867.

Luu, K., Bazin, E., & Blum, M. G. (2017). pcadapt: an R package to perform genome scans for selection based on principal component analysis. Molecular Ecology Resources, 17(1), 67–77.

Marko, P. B., Rogers-Bennett, L., & Dennis, A. B. (2007). MtDNA population structure and gene flow in lingcod (*Ophiodon elongatus*): limited connectivity despite long-lived pelagic larvae. Marine Biology, 150(6), 1301–1311.

Marko, P. B., Hoffman, J. M., Emme, S. A., McGovern, T. M., Keever, C. C., & Nicole Cox, L. (2010). The ‘Expansion–Contraction’ model of Pleistocene biogeography: rocky shores suffer a sea change?. Molecular Ecology, 19(1), 146–169.

Marko, P. B., & Hart, M. W. (2011). The complex analytical landscape of gene flow inference. Trends in Ecology & Evolution, 26(9), 448–456.

Martins, N. T., Macagnan, L. B., Cassano, V., & Gurgel, C. F. D. (2022). Brazilian marine phylogeography: A literature synthesis and analysis of barriers. Molecular Ecology, 31(21), 5423–5439.

MacCready, P., McCabe, R. M., Siedlecki, S. A., Lorenz, M., Giddings, S. N., Bos, J., … & Garnier, S. (2021). Estuarine circulation, mixing, and residence times in the Salish Sea. Journal of Geophysical Research: Oceans, 126(2), e2020JC016738.

Masson, D., & Cummins, P. F. (2000). Fortnightly modulation of the estuarine circulation in Juan de Fuca Strait. Journal of Marine Research, 58(3). https://elischolar.library.yale.edu/journal_of_marine_research/2356

Menge, B., Cerny-Chipman, E., Johnson, A., Sullivan, J., Gravem, S., & Chan, F. (2016). Sea star wasting disease in the keystone predator *Pisaster ochraceus* in Oregon: Insights into differential population impacts, recovery, predation rate, and temperature effects from long-term research. PLOS ONE, 11(5), p.e 0153994.

Miller, K. J., & Ayre, D. J. (2008). Population structure is not a simple function of reproductive mode and larval type: insights from tropical corals. Journal of Animal Ecology, 77(4), 713–724.

Morin, P. A., McCarthy, M. L., Fung, C. W., Durban, J. W., Parsons, K. M., Perrin, W. F., … & Archer, F. I. (2024). Revised taxonomy of eastern North Pacific killer whales (*Orcinus orca*): Bigg’s and resident ecotypes deserve species status. Royal Society Open Science, 11(3), 231368.

Nannan, L., Huamiao, L., Yan, J., Xingan, L., Yang, L., Tianjiao, W., … & Xiumei, X. (2022). Geometric morphology and population genomics provide insights into the adaptive evolution of *Apis cerana* in Changbai Mountain. BMC Genomics, 23(1), 64.

Oulhen, N., Byrne, M., Duffin, P., Gomez-Chiarri, M., Hewson, I., Hodin, J., … & Wares, J. P. (2022). A review of asteroid biology in the context of sea star wasting: possible causes and consequences. The Biological Bulletin, 243(1), 50–75.

Paine, R. T. (1966). Food Web Complexity and Species Diversity. The American Naturalist, 100(910), 65–75. http://www.jstor.org/stable/2459379

Paine, R. T. (1969). A note on trophic complexity and community stability. The American Naturalist, 103(929), 91–93.

Paine, R. T. (1974). Intertidal community structure: experimental studies on the relationship between a dominant competitor and its principal predator. Oecologia, 15, 93–120.

Palumbi, S. R. (1994). Genetic divergence, reproductive isolation, and marine speciation. Annual Review of Ecology and Systematics, 547-572.

Palumbi, S. R. (2004). Marine reserves and ocean neighborhoods: the spatial scale of marine populations and their management. Annual Review of Environment & Resources, 29, 31–68.

Papa, Y., Morrison, M. A., Wellenreuther, M., & Ritchie, P. A. (2022). Genomic stock structure of the marine teleost Tarakihi (*Nemadactylus macropterus*) provides evidence of potential fine-scale adaptation and a temperature-associated Cline amid panmixia. Frontiers in Ecology and Evolution, 10. 10.3389/fevo.2022.862930

Pappalardo, P., Pringle, J. M., Wares, J. P., & Byers, J. E. (2015). The location, strength, and mechanisms behind marine biogeographic boundaries of the east coast of North America. Ecography, 38(7), 722–731.

Pearse, J. S., McClintock, J. B., Vicknair, K. E., & Feder, H. M. (2010). Long-term population changes in sea stars at three contrasting sites. In Echinoderms: Durham. Durham: *Proceedings of the* *12^th^ International Echinoderm Conference* (pp. 633-640).

Petraitis, P. S., & Dudgeon, S. R. (2020). Declines over the last two decades of five intertidal invertebrate species in the western North Atlantic. Communications Biology, 3(1), 591.

Pia, T. S., Johnson, T., & George, S. B. (2012). Salinity-induced morphological changes in *Pisaster ochraceus* (Echinodermata: Asteroidea) larvae. Journal of Plankton Research, 34(7), 590–601.

Pineda, J., Hare, J. A., & Sponaugle, S. U. (2007). Larval transport and dispersal in the coastal ocean and consequences for population connectivity. Oceanography, 20(3), 22–39.

Pinsky, M. L., Selden, R. L., & Kitchel, Z. J. (2020). Climate-driven shifts in marine species ranges: scaling from organisms to communities. Annual Review of Marine Science, 12(1), 153–179.

Popovic, I., Bergeron, L. A., Bozec, Y. M., Waldvogel, A. M., Howitt, S. M., Damjanovic, K., … & Riginos, C. (2024). High germline mutation rates, but not extreme population outbreaks, influence genetic diversity in a keystone coral predator. PloS Genetics, 20(2), e1011129.

Power, M. E., Tilman, D., Estes, J. A., Menge, B. A., Bond, W. J., Mills, L. S., … & Paine, R. T. (1996). Challenges in the quest for keystones: identifying keystone species is difficult—but essential to understanding how loss of species will affect ecosystems. BioScience, 46(8), 609–620.

Pringle, J. M., & Wares, J. P. (2007). Going against the flow: maintenance of alongshore variation in allele frequency in a coastal ocean. Marine Ecology Progress Series, 335, 69–84.

Pringle, J. M., Byers, J. E., Pappalardo, P., Wares, J. P., & Marshall, D. (2014). Circulation constrains the evolution of larval development modes and life histories in the coastal ocean. Ecology, 95(4), 1022–1032.

Raimondi, P. T., Sagarin, R. D., Ambrose, R. F., Bell, C., George, M., Lee, S. F., … & Murray, S. N. (2007). Consistent frequency of color morphs in the sea star *Pisaster ochraceus* (Echinodermata: Asteriidae) across open-coast habitats in the northeastern Pacific. Pacific Science, 61(2), 201–210.

Rambaut, A. (2010). FigTree v1.3.1. *Institute of Evolutionary Biology, University of Edinburgh, Edinburgh*. http://tree.bio.ed.ac.uk/software/figtree/

Riginos, C., & Liggins, L. (2013). Seascape genetics: populations, individuals, and genes marooned and adrift. Geography Compass, 7(3), 197–216.

Robles, C. D., Desharnais, R. A., Garza, C., Donahue, M. J., & Martinez, C. A. (2009). Complex equilibria in the maintenance of boundaries: experiments with mussel beds. Ecology, 90(4), 985–995.

Rogers, R. L., Grizzard, S. L., & Garner, J. T. (2023). Strong, recent selective sweeps reshape genetic diversity in freshwater bivalve *Megalonaias nervosa*. Molecular Biology and Evolution, 40(2), msad024.

Rstudio = Posit team (2024). Rstudio: Integrated Development Environment for R. Posit Software, PBC, Boston, MA. http://www.posit.co/.

Ruiz-Ramos, D., Schiebelhut, L., Hoff, K., Wares, J., & Dawson, M. (2020). An initial comparative genomic autopsy of wasting disease in sea stars. Molecular Ecology, 29(6), 1087–1102. 10.1111/mec.15386

Sanford, E., Roth, M. S., Johns, G. C., Wares, J. P., & Somero, G. N. (2003). Local selection and latitudinal variation in a marine predator-prey interaction. Science, 300(5622), 1135–1137.

Sanford, E., & Menge, B. A. (2007). Reproductive output and consistency of source populations in the sea star *Pisaster ochraceus*. Marine Ecology Progress Series, 349, 1–12.

Sanford, E., & Kelly, M. W. (2011). Local adaptation in marine invertebrates. Annual Review of Marine Science, 3, 509–535.

Sang, H., Li, Y., & Sun, C. (2022). Conservation genomic analysis of the Asian honeybee in China reveals climate factors underlying its population decline. Insects, 13(10), 953.

Schiebelhut, L. M., Giakoumis, M., Castilho, R., Duffin, P. J., Puritz, J. B., Wares, J. P., Wessel, G. M., & Dawson, M. N. (2022a). Minor Genetic Consequences of a Major Mass Mortality: Short-Term Effects in *Pisaster ochraceus*. The Biological Bulletin, 243(3), 328–338. 10.1086/722284

Schiebelhut, L. M., Gaylord, B., Grosberg, R. K., Jurgens, L. J., & Dawson, M. N. (2022b). Species’ attributes predict the relative magnitude of ecological and genetic recovery following mass mortality. Molecular Ecology, 31(22), 5714–5728.

Shanks, A. L. (2009). Pelagic larval duration and dispersal distance revisited. The Biological Bulletin, 216(3), 373–385.

Shapiro, J. S., Chang, H. C., Tatekoshi, Y., Zhao, Z., Waxali, Z. S., Hong, B. J., … & Ardehali, H. (2023). Iron drives anabolic metabolism through active histone demethylation and mTORC1. Nature Cell Biology, 25(10), 1478–1494. 10.1038/s41556-023-01225-6

Sharifian, S., Homaei, A., Kamrani, E., Etzerodt, T., & Patel, S. (2020). New insights on the marine cytochrome P450 enzymes and their biotechnological importance. International Journal of Biological Macromolecules, 142, 811–821.

Shorthouse, D. P. (2010). SimpleMappr, an online tool to produce publication-quality point maps. [Retrieved from https://www.simplemappr.net. Accessed May 31, 2023].

Siegle, M. R., Taylor, E. B., Miller, K. M., Withler, R. E., & Yamanaka, K. L. (2013). Subtle population genetic structure in yelloweye rockfish (*Sebastes ruberrimus*) is consistent with a major oceanographic division in British Columbia, Canada. PloS One, 8(8), e71083.

Silliman, K. (2019). Population structure, genetic connectivity, and adaptation in the Olympia oyster (*Ostrea lurida*) along the west coast of North America. Evolutionary Applications, 12(5), 923–939.

Sobocinski, K.L. (2021). *State of the Salish Sea*. G. Broadhurst and N.J.K. Baloy (Contributing Eds.). Salish Sea Institute, Western Washington University. 10.25710/vfhb-3a69.

Sotka, E. E., Hughes, A. R., Hanley, T. C., & Hays, C. G. (2024). Restricted Dispersal and Phenotypic Response to Water Depth in a Foundation Seagrass. Molecular Ecology, 33(23), e17565.

Stewart-Sinclair, P. J., Last, K. S., Payne, B. L., & Wilding, T. A. (2020). A global assessment of the vulnerability of shellfish aquaculture to climate change and ocean acidification. Ecology and Evolution, 10(7), 3518–3534.

Stickle, W. B., Foltz, D. W., Katoh, M., & Nguyen, H. L. (1992). Genetic structure and mode of reproduction in five species of sea stars (Echinodermata: Asteroidea) from the Alaskan Coast. Canadian Journal of Zoology, 70(9), 1723–1728. 10.1139/z92-239

Strathmann, R. (1978). Length of pelagic period in echinoderms with feeding larvae from the Northeast Pacific. Journal of Experimental Marine Biology and Ecology, 34(1), 23–27.

Sunday, J. M., Bates, A. E., & Dulvy, N. K. (2012). Thermal tolerance and the global redistribution of animals. Nature Climate Change, 2(9), 686–690.

Sutherland, D. A., MacCready, P., Banas, N. S., & Smedstad, L. F. (2011). A model study of the Salish Sea estuarine circulation. Journal of Physical Oceanography, 41(6), 1125–1143.

Therkildsen, N. O., Wilder, A. P., Conover, D. O., Munch, S. B., Baumann, H., and Palumbi, S. R. (2019). Contrasting genomic shifts underlie parallel phenotypic evolution in response to fishing. Science 365:487–490

Thompson, J. D., Higgins, D. G., & Gibson, T. J. (1994). CLUSTAL W: improving the sensitivity of progressive multiple sequence alignment through sequence weighting, position-specific gap penalties and weight matrix choice. Nucleic Acids Research, 22(22), 4673–4680.

Thorson, R. M. (1980). Ice-Sheet Glaciation of the Puget Lowland, Washington, during the Vashon Stade (Late Pleistocene) 1. *Quaternary Research*, *13*(3), 303-321.

Titus, S. E., & Hearther, K. (2019). A disaster for the Pisaster? Temperature effects on ochre sea star larval development. Friday Harbor Laboratories Student Research Papers, 705.

Toczydlowski, R. H., & Waller, D. M. (2019). Drift happens: Molecular genetic diversity and differentiation among populations of jewelweed (Impatiens capensis Meerb.) reflect fragmentation of floodplain forests. Molecular ecology, 28(10), 2459–2475.

Toews, D. P., & Brelsford, A. (2012). The biogeography of mitochondrial and nuclear discordance in animals. Molecular Ecology, 21(16), 3907–3930.

Uthicke, S., Schaffelke, B., & Byrne, M. (2009). A boom–bust phylum? Ecological and evolutionary consequences of density variations in echinoderms. Ecological Monographs, 79(1), 3–24.

Vickery, M. S., & McClintock, J. B. (2000). Effects of food concentration and availability on the incidence of cloning in planktotrophic larvae of the sea star *Pisaster ochraceus*. The Biological Bulletin, 199(3), 298–304.

Villegas-Mirón, P., Acosta, S., Nye, J., Bertranpetit, J., & Laayouni, H. (2021). Chromosome X-wide analysis of positive selection in human populations: common and private signals of selection and its impact on inactivated genes and enhancers. Frontiers in Genetics, 12, 714491.

Wadgymar, S. M., DeMarche, M. L., Josephs, E. B., Sheth, S. N., & Anderson, J. T. (2022). Local adaptation: causal agents of selection and adaptive trait divergence. Annual Review of Ecology, Evolution, and Systematics, 53, 87–111.

Walton, L. N., Watts, V. R., Schuster, J. M., & Bates, A. E. (2024). Cool ocean temperatures fail to buffer the impacts of heat exposure during low tide on the behaviour and physiology of a keystone predator. bioRxiv, 2024-03.

Wang, Q., Liu, Y., Yan, L., Chen, L., & Li, B. (2021). Genome-wide SNP discovery and population genetic analysis of *Mesocentrotus nudus* in China seas. Frontiers in Genetics, 12, 717764.

Wares, J.P., Gaines, S.D., & Cunningham, C.W. (2001). A comparative study of asymmetric migration events across a marine biogeographic boundary. Evolution, 55(2), 295. 10.1111/j.0014-3820.2001.tb01294.x

Wares, J. P., & Pringle, J. M. (2008). Drift by drift: effective population size is limited by advection. BMC Evolutionary Biology, 8(1), 1–7.

Wares, J. P., Strand, A. E., & Sotka, E. E. (2021). Diversity, divergence and density: How habitat and hybrid zone dynamics maintain a genomic cline in an intertidal barnacle. Journal of Biogeography, 48(9), 2174–2185.

Wares, J. P., Duffin, P. Joy (2024) Whole genome sequence data for *Pisaster ochraceus* samples collected from the Pacific coast of North America from July 2004 to May 2018. Biological and Chemical Oceanography Data Management Office (BCO-DMO). (Version 1) Version Date 2024-08-07. doi:10.26008/1912/bco-dmo.934772.1

Whitlock, M. C., & Lotterhos, K. E. (2015). Reliable detection of loci responsible for local adaptation: Inference of a null model through trimming the distribution of FST. The American Naturalist, 186(S1), S24–S36.

Whittaker, K. A., & Rynearson, T. A. (2017). Evidence for environmental and ecological selection in a microbe with no geographic limits to gene flow. Proceedings of the National Academy of Sciences, 114(10), 2651–2656.

Wilder, A. P., Palumbi, S. R., Conover, D. O., & Therkildsen, N. O. (2020). Footprints of local adaptation span hundreds of linked genes in the Atlantic silverside genome. Evolution Letters, 4(5), 430–443.

Williams, P. M., & Chan, K. S. (1966). Distribution and speciation of iron in natural waters: transition from river water to a marine environment, British Columbia, Canada. Journal of the Fisheries Board of Canada, 23(4), 575–593.

Wilson, L. J., Fulton, C. J., Hogg, A. M., Joyce, K. E., Radford, B. T., & Fraser, C. I. (2016). Climate-driven changes to ocean circulation and their inferred impacts on marine dispersal patterns. Global Ecology and Biogeography, 25(8), 923–939.

Wray, A., Petrou, E., Nichols, K. M., Pacunski, R., LeClair, L., Andrews, K. S., … & Hauser, L. (2025). Divergent population structure in five common rockfish species of Puget Sound, WA suggests the need for species-specific management. Molecular Ecology, 34(1), e17590.

Xuereb, A., d’Aloia, C. C., Andrello, M., Bernatchez, L., & Fortin, M. J. (2021). Incorporating putatively neutral and adaptive genomic data into marine conservation planning. Conservation Biology, 35(3), 909–920.

Zakas, C., & Wares, J. P. (2012). Consequences of a poecilogonous life history for genetic structure in coastal populations of the polychaete *Streblospio benedicti*. Molecular Ecology, 21(22), 5447–5460.

Zerebecki, R. A., Sotka, E. E., Hanley, T. C., Bell, K. L., Gehring, C., Nice, C. C., … & Hughes, A. R. (2021). Repeated genetic and adaptive phenotypic divergence across tidal elevation in a foundation plant species. The American Naturalist, 198(5), E152–E169.

Zhang, W., Yang, M., Zhou, M., Wang, Y., Wu, X., Zhang, X., Ding, Y., Zhao, G., Yin, Z., & Wang, C. (2020). Identification of signatures of selection by whole-genome resequencing of a Chinese native pig. Frontiers in Genetics, 11. 10.3389/fgene.2020.566255

## References

Dawson, M. N., Duffin, P. J., Giakoumis, M., Schiebelhut, L. M., Beas-Luna, R., Bosley, K. L.,… & Wares, J. P. (2023). A decade of death and other dynamics: deepening perspectives on the diversity and distribution of sea stars and wasting. The Biological Bulletin, 244(3), 000–000.

## References

Chen, Z., & Narum, S. R. (2021). Whole genome resequencing reveals genomic regions associated with thermal adaptation in redband trout. Molecular Ecology, 30(1), 162–174.

Igoshin, A., Yudin, N., Aitnazarov, R., Yurchenko, A. A., & Larkin, D. M. (2021). Whole-genome resequencing points to candidate DNA loci affecting body temperature under cold stress in Siberian cattle populations. Life, 11(9), 959.

Kon, T., Pei, L., Ichikawa, R., Chen, C., Wang, P., Takemura, I., … & Jiang, L. (2021). Whole-genome resequencing of large yellow croaker (*Larimichthys crocea*) reveals the population structure and signatures of environmental adaptation. Scientific Reports, 11(1), 11235.

Li, M., Tian, S., Yeung, C. K., Meng, X., Tang, Q., Niu, L., … & Li, R. (2014). Whole-genome sequencing of Berkshire (European native pig) provides insights into its origin and domestication. Scientific Reports, 4(1), 4678.

Qiu, Q., Wang, L., Wang, K., Yang, Y., Ma, T., Wang, Z., … & Liu, J. (2015). Yak whole-genome resequencing reveals domestication signatures and prehistoric population expansions. Nature Communications, 6(1), 10283.

Sandercock, A. M., Westbrook, J. W., Zhang, Q., Johnson, H. A., Saielli, T. M., Scrivani, J. A., … & Holliday, J. A. (2022). Frozen in time: Rangewide genomic diversity, structure, and demographic history of relict American chestnut populations. Molecular Ecology, 31(18), 4640–4655.

Zhang, D. Y., Zhang, X. X., Li, F. D., Yuan, L. F., Li, X. L., Zhang, Y. K., … & Wang, W. M. (2022). Whole-genome resequencing reveals molecular imprints of anthropogenic and natural selection in wild and domesticated sheep. Zoological Research, 43(5), 695.

Zheng, J., Zhao, L., Zhao, X., Gao, T., & Song, N. (2022). High genetic connectivity inferred from whole-genome resequencing provides insight into the phylogeographic pattern of *Larimichthys polyactis*. Marine Biotechnology, 24(4), 671–680.

## References

Andrews, S. (2010). FastQC: A quality control tool for high throughput sequence data. Available online at: http://www.bioinformatics.babraham.ac.uk/projects/fastqc/

## References

Danecek, P., Auton, A., Abecasis, G., Albers, C. A., Banks, E., DePristo, M. A., … 1000 Genomes Project Analysis Group. (2011). The variant call format and VCFtools. Bioinformatics, 27(15), 2156–2158.

## References

Altschul, S. F., Gish, W., Miller, W., Myers, E. W., & Lipman, D. J. (1990). Basic local alignment search tool. Journal of Molecular Biology, 215(3), 403–410.

## References

Bosch, S., Fernandez, S. (2023). sdmpredictors: Species Distribution Modelling Predictor Datasets. R package version 0.2.15, http://lifewatch.github.io/sdmpredictors/.

## References

Andrews, K. S., Nichols, K. M., Elz, A., Tolimieri, N., Harvey, C. J., Pacunski, R., … & Tonnes, D. M. (2018). Cooperative research sheds light on population structure and listing status of threatened and endangered rockfish species. Conservation Genetics, 19, 865–878.

Buonaccorsi, V. P., Kimbrell, C. A., Lynn, E. A., & Vetter, R. D. (2002). Population structure of copper rockfish (*Sebastes caurinus*) reflects postglacial colonization and contemporary patterns of larval dispersal. Canadian Journal of Fisheries and Aquatic Sciences, 59(8), 1374–1384. 10.1139/f02-101

COSEWIC. (2008). COSEWIC assessment and status report on the Yelloweye Rockfish *Sebastes ruberrimus*, Pacific Ocean inside waters population and Pacific Ocean outside waters population, in Canada. Committee on the Status of Endangered Wildlife in Canada, 7, 75.

Cunningham, K. M., Canino, M. F., Spies, I. B., & Hauser, L. (2009). Genetic isolation by distance and localized fjord population structure in Pacific cod (*Gadus macrocephalus*): limited effective dispersal in the northeastern Pacific Ocean. Canadian Journal of Fisheries and Aquatic Sciences, 66(1), 153–166. 10.1139/F08-199

Dimond, J. L., Crim, R. N., Unsell, E., Barry, V., & Toft, J. E. (2022). Population genomics of the basket cockle *Clinocardium nuttallii* in the southern Salish Sea: Assessing genetic risks of stock enhancement for a culturally important marine bivalve. Evolutionary Applications, 15(3), 459–470.

Drinan, D. P., Gruenthal, K. M., Canino, M. F., Lowry, D., Fisher, M. C., & Hauser, L. (2018). Population assignment and local adaptation along an isolation-by-distance gradient in Pacific cod (*Gadus macrocephalus*). Evolutionary Applications, 11(8), 1448–1464.

Gregory, R. (1976). Larval dynamics of the Dungeness crab, Cancer magister, off the central Oregon coast, 1970-71. *Fishery Bulletin*, *74*(2).

Hinckley, S., Stockhausen, W. T., Coyle, K. O., Laurel, B. J., Gibson, G. A., Parada, C., … & Ladd, C. (2019). Connectivity between spawning and nursery areas for Pacific cod (*Gadus macrocephalus*) in the Gulf of Alaska. Deep Sea Research Part II: Topical Studies in Oceanography, 165, 113–126.

Hollowed, A. B. (1992). Variability of winter ocean conditions and strong year classes of Northeast Pacific groundfish. In ICES Marine Science Symposium, 195, pp. 433–444.

Iwamoto, E. M., Elz, A. E., García-De León, F. J., Silva-Segundo, C. A., Ford, M. J., Palsson, W. A., & Gustafson, R. G. (2015). Microsatellite DNA analysis of Pacific hake *Merluccius productus* population structure in the Salish Sea. ICES Journal of Marine Science, 72(9), 2720–2731.

Jackson, T. M., & O’Malley, K. G. (2017). Comparing genetic connectivity among Dungeness crab (*Cancer magister*) inhabiting Puget Sound and coastal Washington. Marine Biology, 164, 1–12.

Kendall, A.W. (1989) Additions to knowledge of Sebastes larvae through recent rearing. Alaska Fisheries Science Center, NOAA report 23132_DS1.

Longo, G. C., Head, M. A., Parker-Stetter, S. L., Taylor, I. G., Tuttle, V. J., Billings, A. A., … & Nichols, K. M. (2024). Population genomics of coastal Pacific Hake. North American Journal of Fisheries Management, 44(1), 222–234.

Lowell, N., Suhrbier, A., Tarpey, C., May, S., Carson, H., & Hauser, L. (2023). Population structure and adaptive differentiation in the sea cucumber *Apostichopus californicus* and implications for spatial resource management. PLoS One, 18(3), e0280500.

Moura, A. E., Kenny, J. G., Chaudhuri, R., Hughes, M. A., J. Welch, A., Reisinger, R. R., … Hoelzel, A. R. (2014). Population genomics of the killer whale indicates ecotype evolution in sympatry involving both selection and drift. Molecular Ecology, 23(21), 5179–5192.

Shields, M. W., Hysong-Shimazu, S., Shields, J. C., & Woodruff, J. (2018). Increased presence of mammal-eating killer whales in the Salish Sea with implications for predator-prey dynamics. PeerJ, 6, e6062.

Siegle, M. R., Taylor, E. B., Miller, K. M., Withler, R. E., & Yamanaka, K. L. (2013). Subtle population genetic structure in yelloweye rockfish (*Sebastes ruberrimus*) is consistent with a major oceanographic division in British Columbia, Canada. PLoS One, 8(8), e71083.

Strathmann, R. (1978). Length of pelagic period in echinoderms with feeding larvae from the Northeast Pacific. Journal of Experimental Marine Biology and Ecology, 34(1), 23–27. doi: 10.1016/0022-0981(78)90054-0

